# EpCAM sustains crypt mechanics and prevents aberrant remodeling

**DOI:** 10.64898/2026.04.27.721034

**Authors:** Soleilhac Matis, Ruiz Laura, Berrebi Dominique, Barai Amlan, Lecouffe Brice, Richelme Sylvie, Obino Valentine, Dang Tien, Buzhinsky Nicolas, Rico Felix, Salomon Julie, Ruemmele Frank, Delacour Delphine

## Abstract

Crypt morphogenesis is a fundamental process that drives intestinal growth and maintains its homeostasis. Although intestinal tissue mechanics is recognized as pivotal for early crypt initiation and budding, the molecular mechanisms that couple epithelial forces to mature crypt behavior remain largely unexplored. In particular, while Paneth cells have been proposed to be implicated in the mechanical control of crypt remodeling and fission, the underlying pathways have remained elusive. Here, we identified EpCAM (<u>E</u>pithelial <u>C</u>ell <u>A</u>dhesion <u>M</u>olecule) as a key regulator of epithelial contractility and crypt morphogenesis. EpCAM was preferentially enriched at lateral interfaces of Paneth cells *in vivo* and in organoid models, where it maintained cortical tension and contractile organization. Loss of EpCAM disrupted crypt cell cortical myosin II-A localization, leading to aberrant Paneth cell apical contractility, enlarged apical surfaces, distorted pyramidal morphology, and altered aspect ratios. Consequently, Paneth cells lose their tight clustering in crypt base, their spatial distribution along the crypt axis becomes irregular, and the collective architecture of the crypt epithelium was compromised. At the tissue level, EpCAM deficiency profoundly perturbed crypt fission, increasing fission frequency while favoring asymmetric outcome both in *EPCAM*-mutated Congenital Tufting Enteropathy (CTE) patients. Live imaging of EpCAM-knockout mouse organoids confirmed a failure in the spatial control of crypt bifurcation. Collectively, our findings revealed that EpCAM is not required to initiate fission, but is essential to coordinate the mechanical forces and ensure correct division of the crypt. We showed that EpCAM-dependent biomechanical integrity of Paneth cells generates a stabilizing force field at the crypt base, orchestrating coordinated tissue remodeling. Disruption of this program led to unbalanced force transmission, distorted crypt geometry, and aberrant fission outcomes, revealing a critical link between epithelial mechanics and intestinal morphogenesis.

## INTRODUCTION

Far from being passive structures, tissues are dynamic systems shaped by constant dialogue between biochemical signals and mechanical forces. These physical forces, stemming for instance from cortical tensions or intracellular pressure, do more than just act on cells: they guide essential processes like cell adhesion and cytoskeletal organization. While much attention has traditionally focused on molecular pathways, it is now clear that proper regulation of forces is equally vital in sculpting tissues during development and preserving their integrity throughout life. When this mechanical balance is disrupted, the consequences contribute to development of pathological conditions and even cancer ^1,2,3^.

Epithelial tissues have become a central focus for many biologists and biophysicists, as mechanical forces play a pivotal role in key epithelial developmental processes. In recent years, the intestinal epithelium has emerged as a model of choice for investigating these aspects, largely thanks to the development of *in vitro* mimetic systems such as organoids ^4,5,6,7^. The intestinal tissue is in constant proliferation, making it one of the most rapidly regenerating tissue in adult mammals. The epithelial cell monolayer lining the small intestine displays a structured organization, with proliferative cell in the crypts and differentiated cells in villi ^8,9^. Intestinal stem cells at crypt bases adhere and receive signals from Paneth cells and stroma cells to maintain self-renewal. As cells migrate upward, they lose stemness while still proliferating, and commit to absorptive (enterocytic) or secretory lineages (goblet, enteroendocrine cells) ^8,9,10^.

Work arising from colleagues and our lab demonstrated the requirement of the protein EpCAM (<u>Ep</u>ithelial <u>C</u>ell <u>A</u>dhesion <u>M</u>olecule) in the deployment of tension forces in differentiated epithelial cells ^11,12,13^. Also known as Trop1 or TACSTD1 (Tumor-associated calcium signal transducer 1), EpCAM is a basolateral transmembrane molecule specifically expressed in epithelia, and it is classified as an ‘unconventional cell adhesion molecule’. EpCAM indirectly supports cell adhesion as it modulates junctional architecture through regulation of the actomyosin cytoskeleton ^14,15,11,12,13^. EpCAM also participates in signaling, migration, proliferation and differentiation ^16^. Moreover, EpCAM is directly implicated in the development of a rare childhood enteropathy, the CTE (<u>C</u>ongenital <u>T</u>ufting <u>E</u>nteropathy) also called Intestinal Epithelial Dysplasia (MIM#613217). Non-sense or truncating mutations of *EPCAM*, resulting in the absence of the protein, lead to the development of the disease in 75% of CTE patients ^17,18^. Histologically, the CTE epithelium is defined by distinctive tissue lesions known as ‘tufts’ along the villi. To date, no consistent or characteristic features of hyperplasia or inflammation have been reported in CTE disease ^19,20,21^.

At the molecular level, EpCAM’s role in regulating signaling pathways upstream of myosin-II activity is crucial for epithelial cell organization and tissue architecture. Fagotto and colleagues have shown that EpCAM regulates the level of actomyosin activation via nPKC ^11^. Our team has shown that EpCAM conditions the subcellular dynamics of the activated form of RhoA in the cortex of intestinal epithelial cells. The absence of EpCAM leads to blockage of RhoA-GTP in recycling endosomes dedicated to rapid protein recycling at the cell cortex ^13^. Then, the loss of EpCAM results in actomyosin mislocalization in intestinal cells: instead of homogenously localizing apically beneath the brush border of microvilli, actomyosin develops its activity at multicellular contact points, resulting in deformations of the apical face and tight junction belt, and ultimately in the loss of apical polarity and cellular misalignment of enterocytes. These biomechanical defects are at the root of the characteristic tissue lesions that form along the villi in *EPCAM*-mutated CTE patients ^12,22^.

The role of tissue mechanics in shaping intestinal crypt architecture has begun to receive increasing attention. Crypt formation occurs late during intestinal development. It takes place during gestation in humans (weeks 11-12) but during the first days after birth in mice ^23,10^, and has been recently well-described in the organoid model ^24^. Serra et al. demonstrated that crypt budding is initiated by a symmetry-breaking event mediated through the YAP/TAZ pathway ^24^. This process is notably driven by the emergence of apical constriction forces within the crypt ^25,5^. In addition, Trepat and colleagues revealed, using organoid-derived monolayers, that a contractile mechanical belt forms at the crypt–villus boundary, contributing to the maintenance of crypt compartmentalization ^26^. Once formed, the crypt undergoes tissue remodeling, notably through crypt fission ^23^. However, the role of tissue mechanics in driving crypt fission remains unclear. While pharmacological inflation has been shown to induce crypt branching in organoids, this phenomenon may occur *in vivo* only under pathological conditions ^27^.

In this study, we sought to explore how mechanical regulation influences crypt behavior after its formation. To address this, we employed a candidate-based approach centered on EpCAM, examining its role within the crypt domain.

## RESULTS

### Proper mechanical properties of the crypt rely on EpCAM expression

Although EpCAM has been extensively studied in the literature, its presence in intestinal crypts has been poorly described ^28^. We therefore set out to analyze its distribution along the crypt-villus axis of the intestinal tissue (Figure 1A). Examination of EpCAM immunostaining on human small intestine paraffin sections revealed that, while it is detected in villus epithelial cells as previously described ^12^, this protein is also present in crypt epithelial cells at comparable levels (Figure 1B-C). We therefore hypothesized that it may play a major role in regulating tissue mechanics within crypts. To investigate EpCAM’s participation *in vitro*, we generated inducible EpCAM-knock-out (KO) and non-targeting control-KO murine intestinal organoid lines (Supplementary Figure 1A-B). The EpCAM-KO lineage obtained is a mixed population containing on average 50% EpCAM-negative organoids and 50% EpCAM-positive organoids (Supplementary Figure 1C).

**Figure 1:**
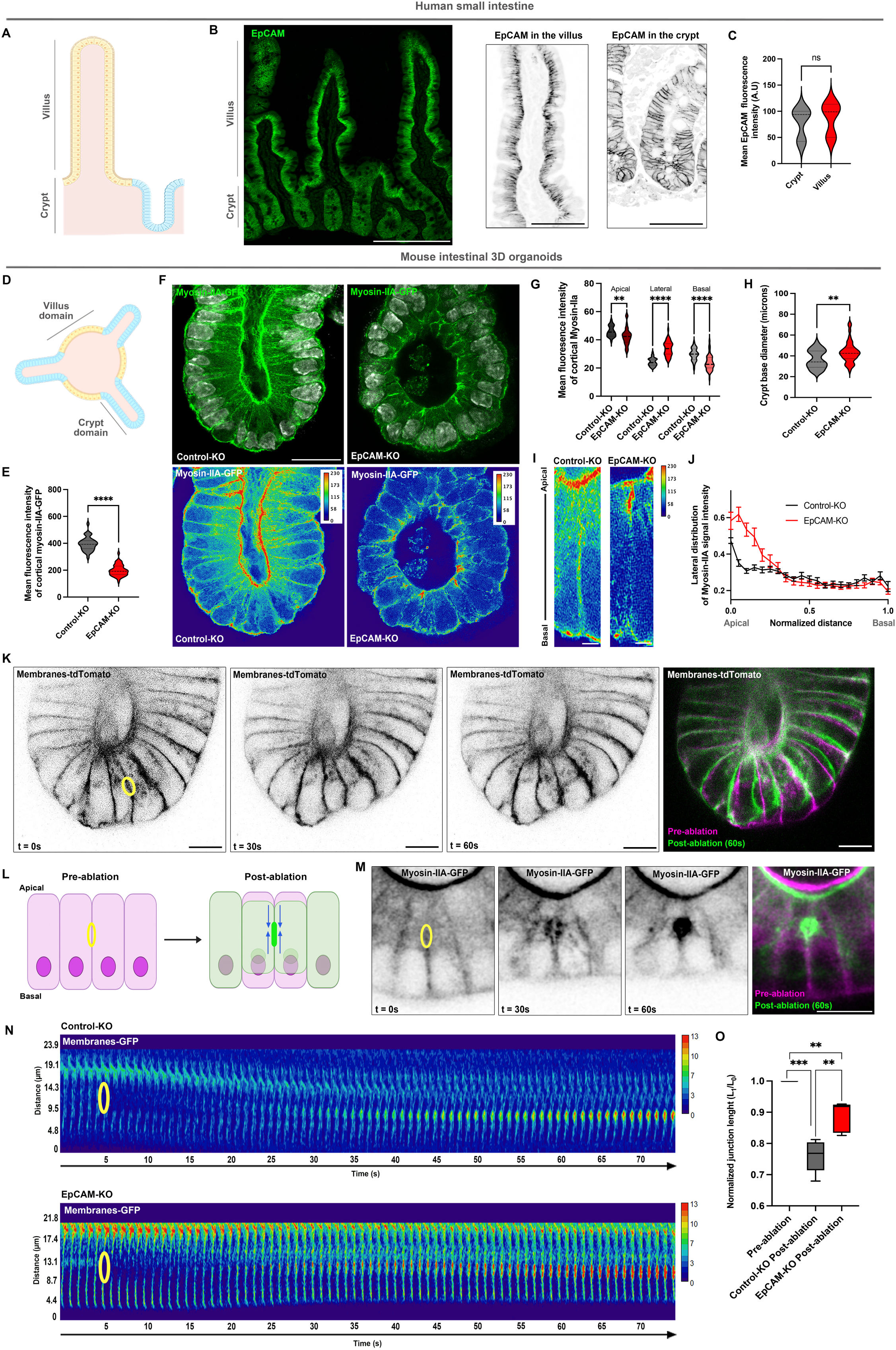
**A,** Scheme showing the functional organization of the mouse small intestinal tissue where the proliferative crypt domain is shown in blue and the differentiated villus compartment in yellow. **B,** Confocal microscopy analysis of the distribution of EpCAM in human small intestine. Scale bar, 200μm. Representative images of EpCAM (grey) localization in the villus and the crypt are shown on the right. Scale bars, 50 μm. **C**, Statistical analysis of EpCAM distribution in villi and crypts in human small intestine. Mean EpCAM signal intensity in the villi = 877.8±91.2 (mean±S.E.M), in the crypts = 769.5±46.20. n (villi) = 6, n (crypts) = 30. Unpaired t-test, ns non-significant. **D,** Scheme showing the organization of the mouse small intestinal organoid where the proliferative crypt-like domains are shown in blue and the differentiated villus-like domains in yellow. **E**, Statistical analysis of the mean cortical myosin-IIA-KI-GFP signal intensity in control-KO or EpCAM-KO organoid crypts. Mean signal intensity in control-KO = 395.5±8.53 (mean±S.E.M), in EpCAM-KO = 198.7±6.23. n (control-KO crypts) = 35, n (EpCAM-KO crypts) = 45. Unpaired t test, **** p < 0.0001. **F**, Confocal microscopy analysis of myosin-IIA-KI-GFP (green) and DNA (grey) in control-KO and EpCAM-KO organoids. Color-coding of myosin-II-A-GFP signal intensity with Physics LUT ImageJ is shown. Calibration bar is added as a right insert. Scale bar, 20μm. **G**, Statistical analysis of the percentage of cortical myosin-IIA-GFP signal intensity in the apical, lateral and basal domains in control-KO or EpCAM-KO organoid crypts. Mean percentage in the apical domain in control-KO = 45.97±0.64% (mean±S.E.M), in EpCAM-KO = 42.43±0.89%, in the lateral domain in control-KO = 24.31±0.48%, in EpCAM-KO = 33.84±0.76%, in the basal domain in control-KO = 29.73±0.72%, in EpCAM-KO = 23.73±0.79%. n (control-KO) = 35 cells, n (EpCAM-KO) = 45 cells. Sidak’s multiple comparisons test, ** p = 0.0036, **** p < 0.0001. **H**, Statistical analysis of the diameter of crypt bases in control-KO and EpCAM-KO organoids. Mean diameter in control-KO = 36.8±1.54 μm (mean±S.E.M), in EpCAM-KO = 43.41±1.52 μm. n (control-KO) = 32 crypts, n (EpCAM-KO) = 43 crypts. Unpaired t test, Welch’s correction, ** p = 0.003. **I**, Distribution of myosin-IIA-KI-GFP signal intensity along lateral contacts in control-KO and EpCAM-KO intestinal organoids. Fluorescence signal intensity is color-coded with Physics LUT ImageJ. Scale bar, 2μm. **J**, Analysis of the apico-basal distribution of myosin-IIA-GFP signal along lateral contacts in control-KO and EpCAM-KO organoids. **K,** Time-lapse images of membranes-tdTomato before or after laser ablation of a lateral contact. The first image (t0) was false-colored in magenta, the last image (t = 60s) was false-colored in green, and images were merged. Scale bar, 5μm. **L**, Scheme depicting the lateral constriction after laser ablation of lateral membranes. Cells before ablation are shown in magenta, after ablation in green. Yellow circle indicted the site of laser ablation. Predicted tensile forces applied there in organoid crypts are shown as blue arrows. **M**, Time-lapse images of myosin-IIA-KI-GFP after laser ablation of a lateral contact. The first image (t0) was false-colored in magenta, the last image (t = 60s) was false-colored in green, and images were merged. Scale bar, 10μm. **N**, Time-lapse images of CellMask Membranes-GFP in control-KO and EpCAM-KO organoids before or after laser ablation of a lateral contact, and presented as kymographs. Laser ablation occurred at t = 5s. Yellow circles point toward the laser ablation site. Signal intensity is color-coded with Physics LUT table from ImageJ. Color scale bar indicates the gray value intensity. **O**, Statistical analyses of the normalized contact length before or after ablation in control-KO or EpCAM-KO organoid crypts. Normalized mean post-ablation contact length in control-KO cells = 0.76±0.02 (mean±S.E.M), in EpCAM-KO cells = 0.89±0.02. n = 5 ablated contacts. Paired t-test for pre-ablation versus control-KO or EpCAM-KO post-ablation, *** p = 0.0005, ** p = 0.0067. Welch’s t-test for control-KO versus EpCAM-KO post-ablation, ** p = 0.0041.

We first assessed the global crypt stiffness by Atomic Force Microscopy (AFM) on 3D intestinal organoid cultures (Supplementary Figure 1D-E). Control crypts exhibited a mean stiffness of 233±10 Pa, whereas EpCAM depletion led to a significant increase in crypt stiffness, reaching 327±15 Pa (Supplementary Figure 1F-G). These data showed that the global rigidity of crypts was impaired by the loss of EpCAM. We next tested the actomyosin distribution in EpCAM-KO organoids. We observed a significant drop in the global fluorescence intensity of cortical myosin-IIA-GFP in EpCAM-KO crypts when compared to control crypts (Figure 1D-F). Furthermore, a notable shift in myosin-IIA-GFP subcellular localization occurred in EpCAM-deficient crypts. The apical pool of cortical myosin-IIA decreased in EpCAM-KO crypts in comparison with controls, directly impacting apical cell constriction and dilatation of the crypts (Figure 1F-H). The basal myosin-IIA pool also decreased in mutant organoids (Figure 1F-H). Moreover, cortical myosin-IIA mislocated along lateral contacts in the absence of EpCAM (Figure 1F-H). There, lateral myosin-IIA–GFP was not homogeneously distributed but exhibited a pronounced accumulation at the most apical region of cell contacts in EpCAM-KO cells (Figure 1F, I-J). These *in vitro* results were in line with data observed *in vivo* in the crypts of CTE biopsies versus control ones, with a redistribution of myosin-IIA signals upon *EPCAM* mutation (Supplementary Figure 1H). Thus, EpCAM silencing altered the distribution of cortical myosin-IIA in crypt cells, potentially impacting their mechanical properties and tensile forces at cell contacts.

To investigate this further, we performed laser ablation in the organoid crypts. No noticeable recoil was observed after laser ablation targeted within crypt cell cytoplasm or at basal membranes (Video 1-2). Remarkably, laser dissection of a lateral contact triggered a constrictive movement in control crypts (Figure 1K-L; Video 3), along with robust post-ablation recruitment of both membranes and myosin-IIA to the laser dissection site (Figure 1M, N; Supplementary Figure 1I-J; Videos 3-4). These data suggested that, contrary to what has been proposed in other model systems ^29^, lateral contacts in the organoid crypt do not generate positive tensile forces but tensile forces orientated in an opposite direction (Figure 1J). Furthermore, in the absence of EpCAM, the magnitude of lateral tensile forces was significantly reduced, as evidenced by the increased normalized contact length following ablation in comparison with controls (from 0.76±0.02 in controls to 0.89±0.02 in EpCAM-KO organoids; Figure 1N-O; Videos 3-4, 6-7). This result was consistent with the apico-lateral accumulation of cortical myosin-IIA and its diminished recruitment along the length of lateral cell-cell contacts in EpCAM-KO crypts (Figure 1F, I-J). Taken together, these results underscored the importance of EpCAM in sustaining the mechanical stability of the intestinal crypt through the regulation of cortical tension.

### Loss of EpCAM leads to abnormal cryptogenesis both in mutant mouse intestinal organoids and in a rare pediatric enteropathy

Based on our observations, we hypothesized that the perturbation of crypt mechanics resulting from EpCAM loss may affect crypt development or structural integrity. We initially assessed EpCAM involvement in cryptogenesis *in vitro* using mouse organoids. No major morphological abnormalities were observed during the early stages of organoid development at day 1 or day 3 in EpCAM-KO organoids in comparison with controls (Supplementary Figure 2A-B). These data indicated that EpCAM is not essential for the initial steps of crypt formation, such as budding and early elongation. However, beginning at day 5 following EpCAM-KO induction, a decrease in the number of crypts per organoid was observed compared to controls (Figure 2A-B; Supplementary Figure 2A-B). In addition, the EpCAM-KO crypts displayed aberrant morphogenesis, characterized by highly elongated, zig-zag tubular structures (Figure 2A,C), and the formation of tissue buds within bent crypt regions (Figure 2A,D). This phenotype persisted through successive passages following EpCAM-KO induction (Supplementary Figure 2A,C), suggesting that the regenerative potential of the stem cell niche itself remained intact despite the loss of EpCAM. To validate the aberrant crypt morphogenesis observed in the absence of EpCAM, we employed intestinal organoids cultured as 2D monolayers, as previously described ^30^, enabling a comprehensive visualization of crypt-villus patterning (Figure 2E). In these organoid monolayers, EpCAM-KO crypt-like domains exhibited a broader and more irregular surface compared to those of control-KO organoids (Figure 2F-G). *In vivo,* cryptogenesis was also defective in *EPCAM*-mutated CTE patients. Alongside the typical villous atrophy ^19,12^, histological analysis of biopsies from CTE patients revealed marked morphogenetic crypt aberrations when compared with those from control patients (Figure 2H-I): crypt surface area (Figure 2H-K), length (Figure 2H-J, L), diameter (Figure 2H-J, M) and lumen surface area (Figure 2H-J, N) were significantly increased under CTE condition in comparison with control biopsies. Moreover, CTE crypts frequently appeared distorted relative to controls, (Figure 2H-I), accompanied by a marked reduction in crypt straightness (straightness mean of 0.96±0.005 (mean±S.E.M) in control crypts and 0.94±0.0007 in CTE crypts; Figure 2H-I, O). Collectively, the *in vitro* findings in murine organoids mirrored the *in vivo* crypt defects observed in human intestinal samples, revealing that EpCAM was essential for proper cryptogenesis.

**Figure 2:**
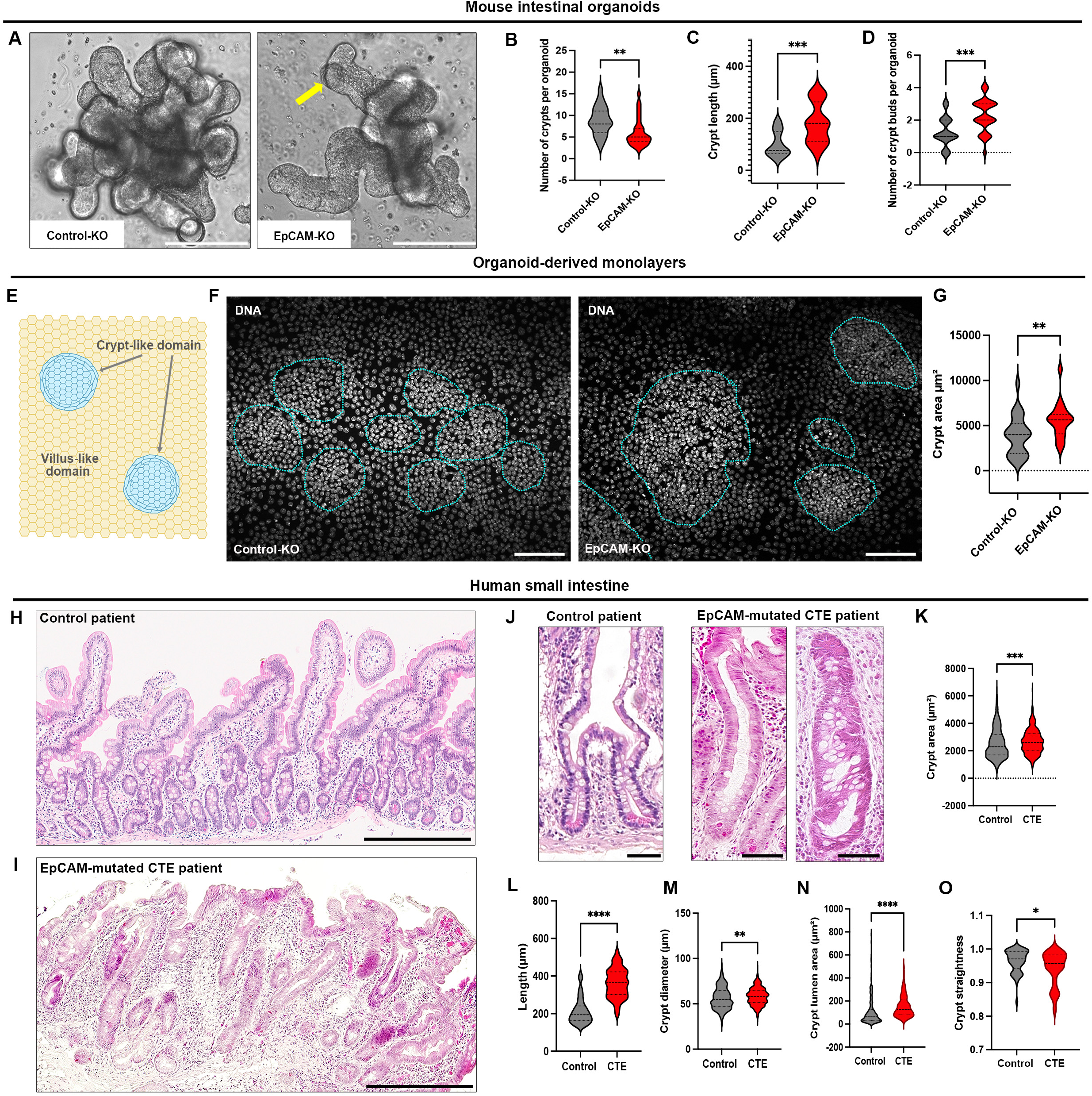
**A,** Brightfield microscopy analysis of the morphology of control-KO and EpCAM-KO organoids after 5 days of culture. Yellow arrow points to a tissue bud in crypt. Scale bar, 200μm. **B**, Statistical analysis of the number of crypts in control-KO and EpCAM-KO organoids. Mean number of crypts in control-KO = 8.83±0.68 crypts (mean±S.E.M), in EpCAM-KO = 6.03±0.58 crypts. Mann-Whitney test, **p = 0.0014. n (control-KO) = 23 organoids, n (EpCAM-KO) = 31 organoids. **C**, Statistical analysis of the crypt length in control-KO and EpCAM-KO organoids. Mean crypt length in control-KO = 101±10.27μm (mean±S.E.M), in EpCAM-KO = 180.7±18.17. Mann-Whitney test, ***p = 0.0006. n (control-KO) = 22 crypts, n (EpCAM-KO) = 18 crypts. **D**, Statistical analysis of the number of buds in crypts of control-KO and EpCAM-KO organoids. Mean number of buds along crypts in control-KO = 1.28±0.16 (mean±S.E.M), in EpCAM-KO = 2.29±0.18. Unpaired t test, ***p = 0.0002. n (control-KO) = 29 organoids, n (EpCAM-KO) = 31 organoids. **E**, Scheme showing the organization of intestinal organoid-derived monolayer with the crypt-like domains in blue and the villus-like compartment in yellow. **F**, Confocal microscopy analysis of nuclei (DNA, gray) in control-KO and EpCAM-KO organoids. Blue dotted lines delimit the crypt-like compartments. Scale bar, 50μm. **G**, Statistical analysis of the crypt area in control-KO and EpCAM-KO organoid-derived monolayers. Mean crypt area in control-KO = 3884±434.1μm^2^ (mean±S.E.M), in EpCAM-KO = 5558±447μm^2^. Mann Whitney test, **p = 0.006. n (control-KO) = 25 crypts, n (EpCAM-KO) = 19 crypts. **H-I,** Histological analysis of hematoxylin/eosin-stained paraffin sections of duodenal biopsies from control (**H**) and EpCAM-mutated CTE (**I**) patients. Scale bar, 150μm**. J,** Representative hematoxylin/eosin-stained duodenal crypts in control and EpCAM-mutated CTE patients. Scale bar, 50μm**. K,** Statistical analysis of crypt area in control and EpCAM-mutated CTE patients. Mean crypt area in control biopsies = 2542±50.1 μm (mean±S.E.M), in CTE biopsies = 2669±40.7 μm. n (control) = 470 crypts, n (CTE) = 458 crypts. Mann-Whitney test, *** p = 0.001. **L**, Statistical analysis of crypt length in control and EpCAM-mutated CTE patients. Mean crypt length in control biopsies = 213.3±8.6 μm (mean±S.E.M), in CTE biopsies = 361.9±6.25 μm. n (control) = 65 crypts, n (CTE) = 154 crypts. Mann-Whitney test, **** p< 0.0001. **M**, Statistical analysis of crypt diameter in control and EpCAM-mutated CTE patients. Mean crypt diameter in control biopsies = 56.39±0.55μm (mean±S.E.M), in CTE biopsies = 58.22±0.45μm. n (control) = 470 crypts, n (CTE) = 458 crypts. Mann-Whitney test, ** p = 0.0013. **N**, Statistical analysis of the area of the crypt lumen in control and EpCAM-mutated CTE patients. Mean lumen area in control biopsies = 110.4±8.18μm^2^ (mean±S.E.M), in CTE biopsies = 157.7±7.36μm^2^. n (control) = 229 crypts, n (CTE) = 182 crypts. Mann-Whitney test, **** p< 0.0001. **O**, Statistical analysis of the crypt straightness in control and EpCAM-mutated CTE patients. Mean straightness in control biopsies = 0.96±0.005 (mean±S.E.M), in CTE biopsies = 0.94±0.0007. n (control) = 51 crypts, n (CTE) = 55 crypts. Mann-Whitney test, * p<0.05.

Next, we characterized the crypt phenotype in the absence of EpCAM. Despite the morphological abnormalities, compartmentalization of the crypt-villus axis and intestinal cell differentiation were preserved. Proliferative cells, marked by EdU or Ki67, remained in the crypt compartment, while CK20^+^-enterocytic differentiated cells were confined to the villus compartment in *EPCAM*-mutated CTE patients (Supplementary Figure 3A-B). *In vitro,* detailed analysis of the cellular composition of EpCAM-KO crypts in organoid-derived monolayers and 3D EpCAM-KO organoids confirmed the presence of Ki67^+^- or EdU^+^-proliferative cells in crypts (Supplementary Figure 3C-D,F-G), along with the exclusion of CK20^+^-differentiated cells (Supplementary Figure 3E,H), typically found in villus-like regions. These data confirmed that EpCAM-KO organoids generated crypt-villus axis, albeit crypts were aberrantly formed. We further evaluated the impact of EpCAM loss on the organization of the stem cell niche. In CTE patient biopsies, no significant difference was observed in the proportion or distribution of OLFM4^+^ intestinal stem cells compared to controls (Supplementary Figure 4A). Similarly, in 3D EpCAM-KO organoids, OLFM4^+^-cells remained enriched at the base of crypts and budding regions (Supplementary Figure 4B). Thus, while EpCAM deficiency disrupted normal crypt maturation, global crypt organization, including tissue compartmentalization and stem cell positioning, remained intact.

### Loss of EpCAM impacts the behavior of Paneth cells and their mechanical homeostasis

To gain deeper insight into the crypt phenotypes under EpCAM deficiency, we examined its cellular distribution within the crypt. *In vivo*, EpCAM signal intensity was detected at lateral cell contacts of crypt cells, but significantly elevated at cell interfaces of lysozyme^+^-Paneth cells compared to other crypt cell interfaces (Figure 3A-B). A similar pattern was observed in organoid-derived monolayers, where EpCAM was concentrated at the periphery of Paneth cells in crypt-like domains (Figure 3C-E). These data revealed a preferential association of EpCAM with Paneth cells in the crypt, suggesting a critical role in regulating Paneth cell mechanics. Cortical myosin-IIA distribution was thus specifically analyzed in this cell type. Paneth cells exhibited higher level of cortical myosin-IIA than other crypt cells (Figure 3F,G control-KO). Similar observation was made *in vivo* in mouse jejunum (Supplementary Figure 5A). Moreover, cortical myosin-IIA distribution was different between Paneth cells and other crypt cells, with more elevated contractility at lateral cortex than apical cortex in Paneth cells (Figure 3H). In the absence of EpCAM, a marked reduction in the proportion of cortical myosin-IIA occurred when compared to control Paneth cells (Figure 3G). This contractile remodeling was particularly perturbed at lateral Paneth cell contacts, with a strong myosin-IIA accumulation at their most apical part (Figure 3I-K). Thus, EpCAM is required for correct contractile patterning in Paneth cells.

**Figure 3:**
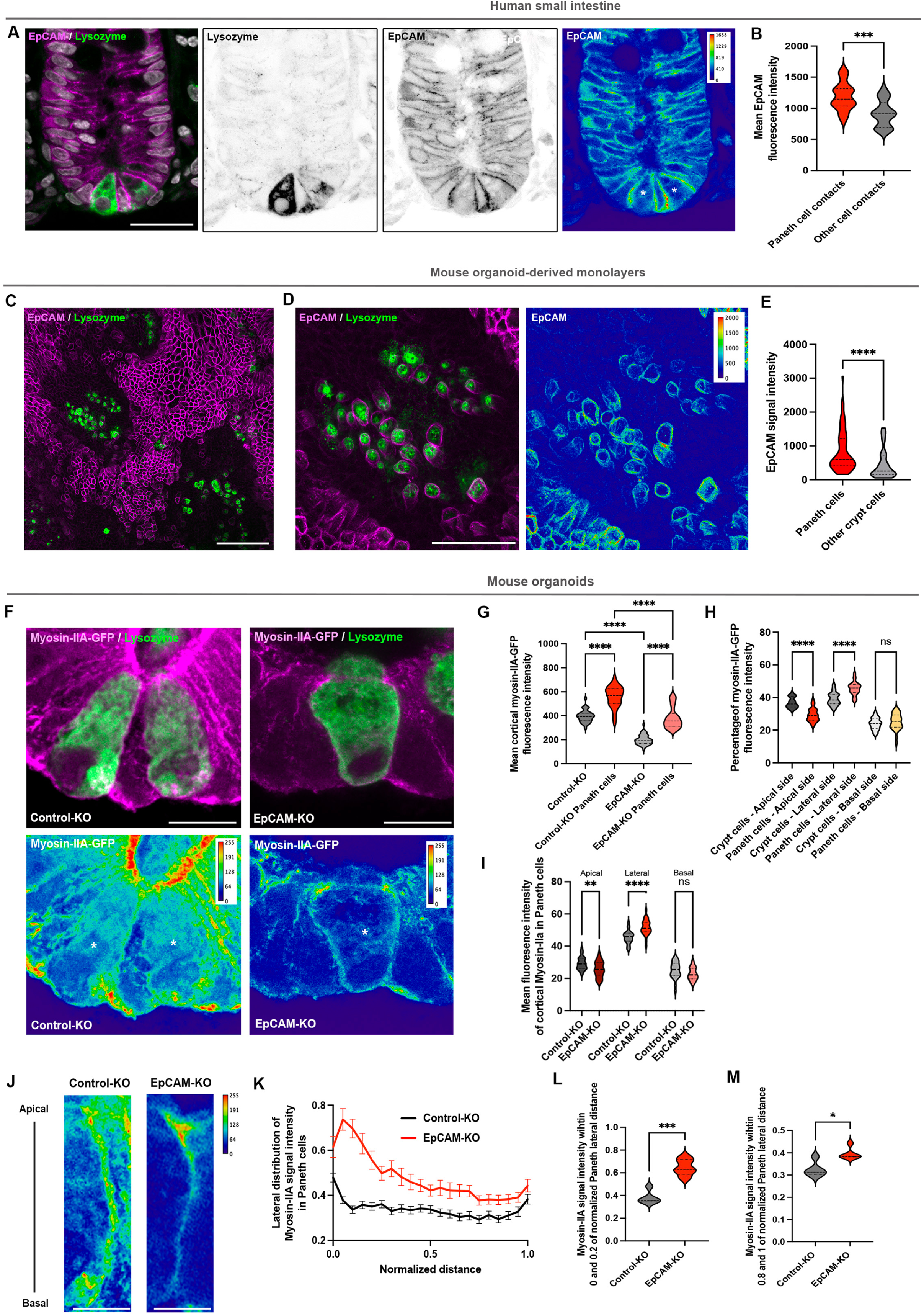
**A**, Confocal microscopy analysis of the distribution of EpCAM (magenta) and lysozyme (green) in crypts of human small intestine. EpCAM signal intensity is color-coded with Physics LUT (lookup table) from ImageJ on the right. Color scale bar indicates the gray value intensity. White stars point to Paneth cells. Scale bar, 20μm. **B**, Statistical analysis of signal intensity level of EpCAM at Paneth cell contacts and other crypt cells in human small intestine. Mean EpCAM signal intensity at Paneth cell contacts = 1190±49.52, at other crypt cell contacts = 916.4±45.15 (mean±S.E.M). n (Paneth cell contacts) = 20 cells, n (other crypt cell contacts) = 22 cells. Unpaired t-test, ***p = 0.0002. **C**, Confocal microscopy analysis of the distribution of EpCAM (magenta) and lysozyme (green) in mouse intestinal organoid monolayers. Scale bar, 100μm. **D**, Confocal microscopy analysis of the distribution of EpCAM (magenta) and lysozyme (green) in crypt-like domains in mouse intestinal organoid monolayers. EpCAM signal intensity is color-coded with Physics LUT from ImageJ on the right. Color scale bar indicates the gray value intensity. Scale bar, 50μm. **E**, Statistical analysis of EpCAM signal intensity at Paneth cell contacts versus at other crypt cell contacts in organoid-derived monolayers. Mean EpCAM signal intensity at Paneth cell contacts = 836.8±75.67 (mean±S.E.M), in other crypt cell contacts = 462.2±63.97. n (Paneth cells) = 65 cells, n (other crypt cells) = 46 cells. Mann-Whitney test, **** p < 0.0001. **F,** Confocal microscopy analysis of myosin-IIA-KI-GFP (magenta) and lysozyme (green) in 3D intestinal organoids. Zoom on a representative Paneth cell is shown on the right. Color-coding of myosin-II signal intensity with Physics LUT ImageJ, and color scale bar indicating the gray value intensity are shown. White star points to Paneth cells. Scale bar, 10μm. **G**, Statistical analysis of the mean cortical myosin-IIA-KI-GFP signal intensity in Paneth cells and other crypt cells in control-KO or EpCAM-KO organoids. Mean cortical signal intensity in control-KO crypt cells = 395.5±8.53 (mean±S.E.M), in control-KO Paneth cells = 558±17.38, in EpCAM-KO crypt cells = 198.7±6.23, in EpCAM-KO Paneth cells = 382.6±16.14. n (control-KO crypt cells) = 35 cells, n (control-KO Paneth cells) = 26 cells, n (EpCAM-KO crypt cells) = 45 cells, n (EpCAM-KO Paneth cells) = 30 cells. One-way ANOVA, **** p < 0.0001. **H**, Statistical analysis of the percentage of the distribution of mean cortical myosin-IIA-KI-GFP signal intensity in Paneth cells and other crypt cells in control-KO organoids. Mean percentage in control-KO organoids in apical cortex of crypt cells = 36.80±0.55% (mean±S.E.M), in apical cortex of Paneth cells = 29.37±0.8%, in lateral cortex of crypt cells = 39.21±0.67%, in lateral cortex of Paneth cells = 45.57±0.8%, in basal cortex of crypt cells = 23.99±0.60%, in basal cortex of Paneth cells = 25.06±1%. n (crypt cells) = 35 cells, n (Paneth cells) = 26 cells. One-way ANOVA, Tukey’s multiple test, **** p < 0.0001. **I**, Statistical analysis of the percentage of the mean cortical myosin-IIA-KI-GFP signal intensity in Paneth cells in contro-KO and EpCAM-KO. Mean percentage in the apical cortex in control-KO Paneth cells = 29.37±0.8%, in EpCAM-KO Paneth cells = 25.66±0.9%, at the lateral cortex in control-KO Paneth cells= 45.57±0.8%, in EpCAM-KO Paneth cells = 51.566±0.86%, at the basal cortex in control-KO Paneth cells = 25.06±1%, in EpCAM-KO Paneth cells = 22.77±0.71%. n (control-KO) = 26 cells, n (EpCAM-KO) = 30 cells. Two-way ANOVA, Sidak’s test, ** p = 0.007, **** p < 0.0001. **J,** Color-coding with Physics LUT ImageJ of representative myosin-IIA-GFP signal intensity at lateral contacts of Paneth cells in control-KO and EpCAM-KO organoids. Color scale bar indicating the gray value intensity are shown. Scale bar, 5μm. **K**, Analysis of the distribution of myosin-IIA-GFP signal intensity along lateral contacts in control-KO and EpCAM-KO Paneth cells. **L,** Statistical analysis of the signal intensity of myosin-IIA-GFP at the most apical part of the Paneth cell lateral contact (between 0 and 0.2 of normalized distance from the apical cell side). Mean intensity in control-KO = 0.38±0.02 (mean± SEM), in EpCAM-KO = 0.65±0.03. Welch’s test, *** p = 0.0002. n (control-KO) = 26 contacts, n (EpCAM-KO) = 29 contacts. **M,** Statistical analysis of the signal intensity of myosin-IIA-GFP at the most basal part of the Paneth cell lateral contact (between 0.8 and 1 of normalized distance from the apical cell side). Mean intensity in control-KO = 0.32±0.01 (mean± SEM), in EpCAM-KO = 0.39±0.01. Mann Whitney’s test, * p = 0.03. n (control-KO) = 26 contacts, n (EpCAM-KO) = 29 contacts.

The consequences of disrupted mechanical homeostasis in EpCAM-mutated Paneth cells were multifaceted. At the cellular level, Paneth cells from CTE patients lost their characteristic pyramidal morphology and displayed an enlarged apical surface when compared with Paneth cells in control patients (Figure 4A). Similar morphological defects were observed *in vitro* in EpCAM-KO organoids (Figure 4B), characterized by a pronounced and selective expansion of the apical domain in EpCAM-KO Paneth cells compared with both other EpCAM-KO crypt cells and control Paneth cells (Figure 4C). In addition, basal lengths were increased in both Paneth and other crypt cells in the absence of EpCAM, with very little variation of cell heights (Figure 4 C; Supplementary Figure 6A). As a result, we defined a cell shape index (ratio between apical area and basal area) of Paneth cells, which was markedly altered for Paneth cells in EpCAM-KO organoids (Figure 4D-E). Paneth cells and other crypt cells exhibited comparable and homogeneous distribution of cell shape indexes in control organoids, with the majority of cell shape indexes comprised between 0 and 0.6 in control-KO crypts (Figure 4F). However, cell shape indexes in EpCAM-KO organoids exhibited marked heterogeneity, particularly among Paneth cells, for which 34.43% were above 1 (Figure 4G, Supplementary Figure 6B-C). Consequently, these cellular alterations in EpCAM-KO organoids may affect the spatial and collective behavior of cells within crypts. To test this hypothesis, we estimated the global cell organization by assessing the arrangement of nuclei in organoid crypts. We thus estimated the 3D orientation of neighboring nuclei by measuring pairwise vector alignment using the dot product (Figure 4H, Supplementary Figure 6D-J). While in control-KO crypts, most of the nuclei were arranged so that they are parallel to one another (parallel orientation, with a normalized dot product between 0.8 and 1), nuclei collectively organized mainly in a non-parallel manner (between 0 and 0.79) in EpCAM-KO crypts (Figure 4H-J). In addition, we analyzed the amplitude and orientation of cell tilting in crypts by measuring the angle formed between each nuclear long axis and the crypt long axis. As previously reported by Perez-Gonzalez et al. ^26^, nuclei displayed a regular and homogenous tilt towards crypt base in control-KO crypts, with the majority of nucleus/crypt angles comprised between 60° and 90°, Figure 4K-L). In EpCAM-KO organoids, nucleus/crypt angles mainly varied between 90° and 120°, revealing that cell tilting displayed markedly greater heterogeneity and crypt cells lost their collective, polarized orientation towards crypt base, when compared with controls (Figure 4K-L). Thus, EpCAM’s loss led to an irregular cell arrangement within the crypt monolayer.

**Figure 4:**
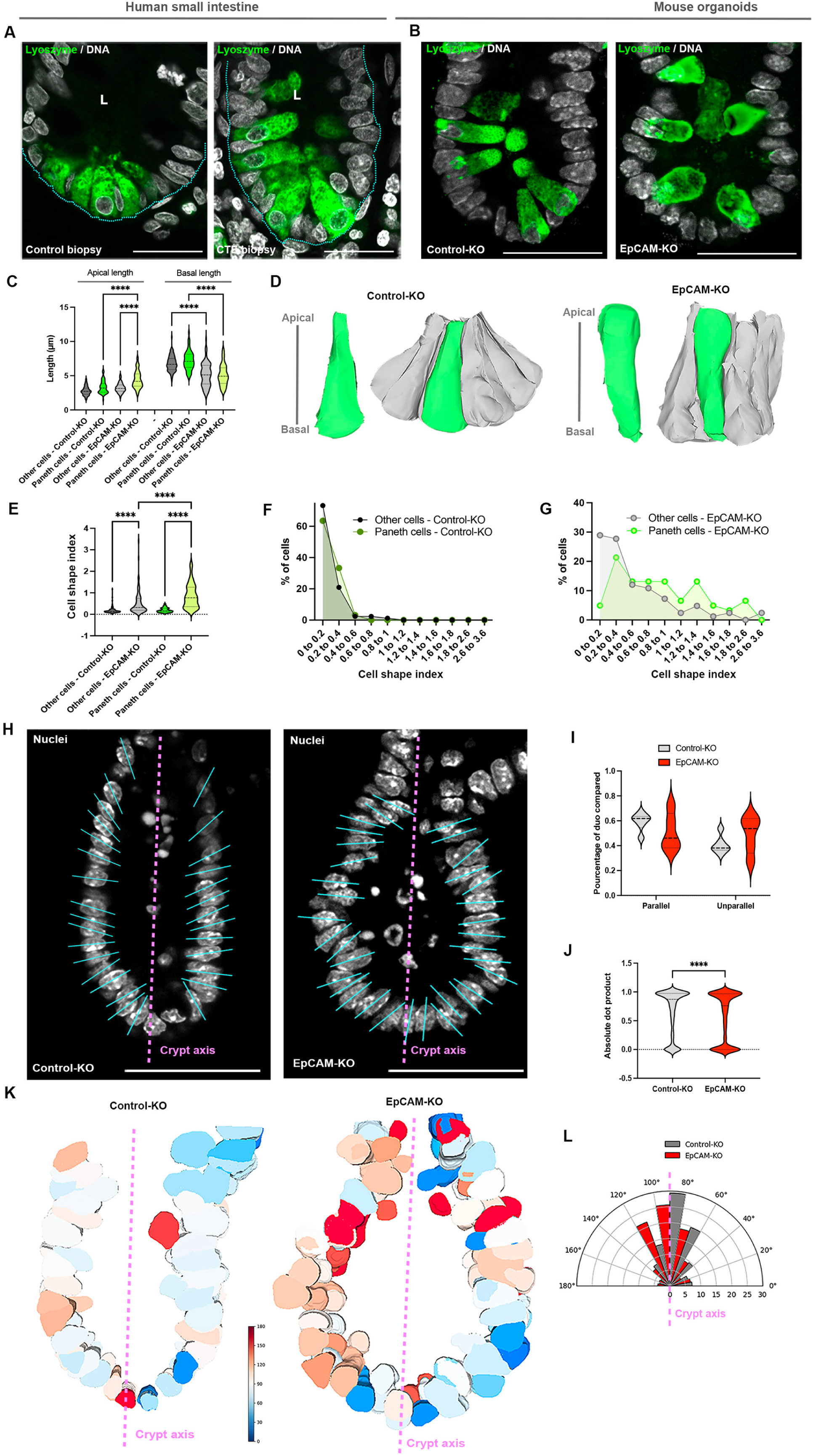
**A**, Confocal microscopy analysis of Paneth cell shape in control or EpCAM-mutated CTE biopsies. Lysozyme-positive cells (green) and nuclei (grey) are shown. Scale bar, 10μm. **B**, Confocal microscopy analysis of Paneth cell shape in control-KO or EpCAM-KO 3D organoids. Lysozyme (green) and DNA (grey) are shown. Scale bar, 20μm**. C,** Statistical analysis of the apical and basal surface lengths of Paneth cells and other crypt cells in control-KO and EpCAM-KO organoids. Mean apical surface length of control-KO other crypt cells = 2.783±0.069 μm (mean±S.E.M), control-KO Paneth cells = 3.334±0.114, EpCAM-KO other crypt cells = 3.229±0.075, EpCAM-KO Paneth cells = 4.432±0.162. Mean basal surface length of control-KO crypt cells = 6.796±0.119 μm (mean±S.E.M), control-KO Paneth cells = 7.203±0.172, EpCAM-KO crypt cells = 5.172±0.186, EpCAM-KO Paneth cells = 5.125±0.198. n (control-KO crypt cells) = 86 cells, n (control-KO Paneth cells) = 63 cells, n (EpCAM-KO crypt cells) = 84 cells, n (EpCAM-KO Paneth cells) = 61 cells. Two-way ANOVA, Tukey’s multiple comparison test, **** p < 0.0001. **D**, Representative 3D rendering of a Paneth cell (green) and its neighbors (grey) after segmentation of cell membranes based on confocal z-stacks in control-KO and EpCAM-KO crypt organoids. **E**, Statistical analysis of cell shape index (apical area/basal area) in control-KO or EpCAM-KO 3D organoid crypts. Mean shape index of other cells in control-KO = 0.187±0.017 (mean±S.E.M), of other cells in EpCAM-KO = 0.534±0.061, of Paneth cells in control-KO = 0.194±0.013, of Paneth cells in EpCAM-KO = 0.859±0.072. n (other cells control-KO) = 86 cells, n (other cells EpCAM-KO) = 83 cells, n (Paneth cells control-KO) = 63 cells, n (Paneth cells EpCAM-KO) = 61 cells. One-way Anova test, Tukey’s multiple comparison test, ****p < 0.0001. **F**, Distribution of cell shape index of Paneth cells (green) and other crypt cells (grey) in control-KO crypts. Cell shape indexes from 0 to 0.2: other cells = 73.25%, Paneth cells = 63.5%, from 0.2 to 0.4: other cells = 20.93%, Paneth cells = 33.33%, from 0.4 to 0.6: other cells = 2.33%, Paneth cells = 3.17%, from 0.6 to 0.8: other cells = 2.33%, Paneth cells = 0%, from 0.8 to 1: other cells = 1.16%, Paneth cells = 0%. **G**, Distribution of cell shape indexes of Paneth cells (green) and other crypt cells (grey) in EpCAM-KO crypts. Cell shape indexes from 0 to 0.2: other cells = 28.91%, Paneth cells = 4.92%, from 0.2 to 0.4: other cells = 27.71%, Paneth cells = 21.32%, from 0.4 to 0.6: other cells = 12.05%, Paneth cells = 13.11%, from 0.6 to 0.8: other cells = 10.84%, Paneth cells = 13.11%, from 0.8 to 1: other cells = 7.24%, Paneth cells = 13.11%, from 1 to 3.6: other cells = 13.25%, Paneth cells = 34.43%. **H**, Confocal analysis of nucleus (grey) distribution in crypts of control-KO and EpCAM-KO organoids. Magenta dotted line shows the crypt axis. Blue lines highlight nucleus long axes. Scale bar, 50μm. **I**, Percentage of neighboring nuclei having a parallel vector orientation (normalized dot product between 0.8 and 1) or non-parallel orientation (between 0 and 0.79). Mean percentage of normalized dot product between 0.8 and 1 in control-KO = 0.594±0.03 (mean±S.E.M), between 0 and 0.79 in control-KO = 0.406±0.03, between 0.8 and 1 in EpCAM-KO = 0.509±0.07, between 0 and 0.79 in EpCAM-KO = 0.49±0.07. **J**, Global statistical analysis of dot products as a proxy of vector orientation of neighboring nuclei in control-KO and EpCAM-KO crypts. Mean dot product in control-KO = 0.663±0.006 (mean±S.E.M), in EpCAM-KO = 0.569±0.007. Welch’s test, ****p < 0.0001. **K**, Representative 3D rendering of nuclear arrangement in control-KO and EpCAM-KO crypt organoids. Nuclear angular deviation is color-coded and color scale bar is shown on the right. **L**, Distribution of the angular deviation between the crypt axis and the nucleus long axis in control-KO and EpCAM-KO organoids.

At the overall tissue level, the distribution of Paneth cells was impacted by EpCAM’s deficiency. *In vivo,* in CTE patients carrying *EPCAM* mutations, Paneth cells were less clustered in crypt base (Figure 5A-B). The average distance between neighboring Paneth cells increased from 11.39 ± 0.38 μm (mean±S.E.M) in control biopsies to 16.01 ± 0.55 μm in CTE patient samples (Figure 5C). Moreover, some Paneth cells in CTE biopsies were scattered along the elongated crypt trunks, with the average distance from the crypt base rising to 36.41 ± 3.08 μm (mean±S.E.M), compared to 14.59 ± 2.62 μm in controls (Figure 5B,D). In 3D EpCAM-KO organoids, this mislocalization of Paneth cells in crypts was even more pronounced along the crypt axis, in contrast to their tight clustering in crypt bottoms of control organoids (Figure 5E-F). A similar observation was made in the organoid-derived monolayers, where Paneth cells were significantly less concentrated in the crypt center (Supplementary Figure 7A-C): the average distance between Paneth cells and the crypt center increased from 24.85 ± 1.30 μm in controls to 34.23 ± 1.81 μm in EpCAM-KO monolayers (Supplementary Figure 7A-C). Altogether, these findings revealed a tissue-level disorder of intestinal crypts arising from the loss of EpCAM and mechanical homeostasis, marked by prominent changes in Paneth cell morphology and behavior within the crypt.

**Figure 5:**
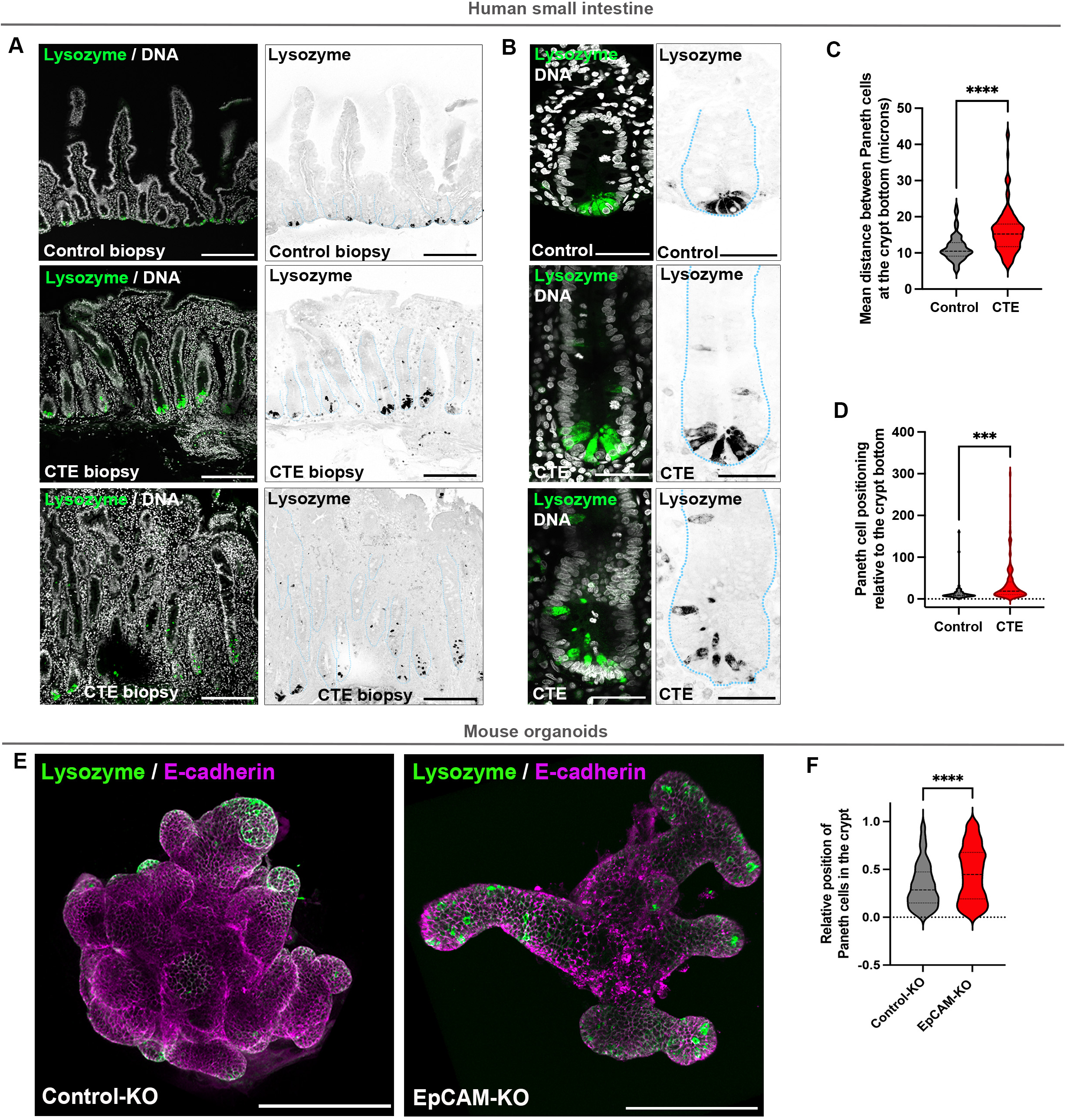
**A-B**, Confocal analysis of Paneth cell distribution in control and EpCAM-mutated CTE patients. Lysozyme (green) and DNA (grey) are shown. Crypts are delimited with a blue dotted line. Scale bars, A 200μm, B 100μm**. C**, Statistical analysis of the mean distance between Paneth cell nucleus centroids in crypts of control and EpCAM-mutated CTE patients. Mean distance in control crypts = 11.39±0.38μm (mean±S.E.M), in CTE crypts = 16.01±0.55μm. Unpaired t test, **** p < 0.0001. N = 6 control biopsies, N = 7 CTE biopsies, n (control) = 87 Paneth cells, n (CTE) = 127 Paneth cells. **D**, Statistical analysis of the distribution of Paneth cells in crypts of control and EpCAM-mutated CTE patients. Mean distance between a Paneth cell and the crypt bottom in control biopsies = 14.59±2.62μm (mean±S.E.M), in CTE biopsies = 36.41±3.08μm. Unpaired t test, ***p<0.0001. N = 5 control patients, N = 5 CTE patients. n (control) = 9 crypts, n (CTE) = 31 crypts. **E**, Confocal microscopy analysis of lysozyme (green) and E-cadherin (magenta) in control-KO and EpCAM-KO 3D organoids. Scale bar, 200μm. **F**, Statistical analysis of the relative position of Paneth cells in crypts of control-KO and EpCAM-KO organoids. Mean relative position of Paneth cells in control-KO organoids = 0.333±0.016 (mean±S.E.M), in EpCAM-KO = 0.453±0.012. Mann-Whitney test, ****p<0.0001. n (Paneth cells in control-KO) = 203 cells, n (Paneth cells in EpCAM-KO) = 537 cells.

### EpCAM guarantees the proper progression of crypts through the fission process

Paneth cells constitute a central structural and functional component of the intestinal stem cell niche, ensuring stem cell maintenance through both biochemical signaling at cell–cell interfaces and intimate physical interactions that compact and organize the crypt architecture ^31,32^. Beyond these canonical roles, pioneering work from I. Näthke’s group has proposed that Paneth cells may shape tissue geometry through force generation and transmission, thereby acting as mechanical regulators of intestinal crypt fission ^33^. Once established, intestinal crypts undergo architectural remodeling through crypt fission, a process that is indispensable for postnatal intestinal growth and the maintenance of adult tissue homeostasis ^34,35^. Crypt fission is viewed as a slow, cyclical morphogenetic program initiated at the crypt base, in which the mother crypt undergoes progressive enlargement, followed by the emergence of a nascent tissue bud that ultimately bifurcates into two daughter crypts (D1-D2; Figure 6A) ^33,36^. Literature reports that disturbances during crypt cycle leads to asymmetrical fission and ‘corrupted’ crypts (Figure 6B; ^33,37,38^. Here, our data revealed profound defects in crypt contractility accompanied by alterations in the mechanical properties of Paneth cells upon loss of EpCAM. These observations lead us to advance the hypothesis that EpCAM-dependent mechanical integrity of Paneth cells may be a critical driver of crypt fission, and that its disruption may underlie the severe crypt architectural phenotypes observed in EpCAM-deficient tissues.

**Figure 6:**
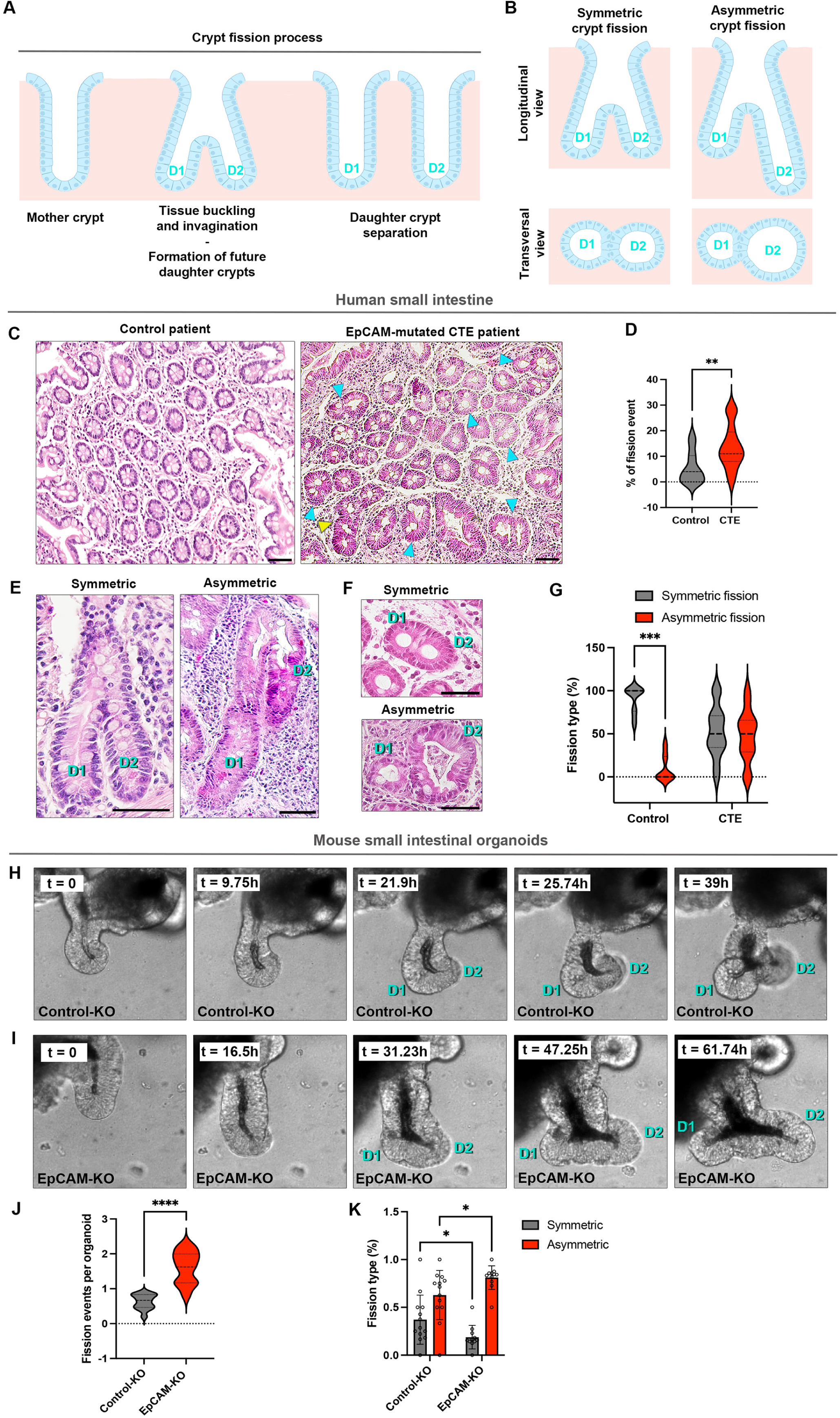
**A,** Schematics showing the morphogenetic steps described in literature for crypt fission and generation of two daughter crypts (D1-D2). **B**, Schematics depicting the crypt morphology during symmetric aor asymmetric fission. **C,** Histological analysis of hematoxylin/eosin-stained paraffin sections of duodenal biopsies from control and EpCAM-mutated CTE patients. Blue arrowheads point to asymmetric crypt fission. Yellow arrowheads point to multifide crypt fission. Scale bar, 50μm. **D**, Statistical analysis of the number of crypt fission events in control and EpCAM-mutated CTE patients. Mean number of crypt fission events in control biopsies = 7.16±1.65 % (mean±S.E.M), in CTE biopsies= 14.31±1.50%. Mann Whitney test, ** p = 0.0039. n (control) = 12 patients, n = (CTE) = 13 patients. **E-F**, Histological analysis of daughter crypt shapes in longitudinal (**E**) or transversal views (**F**) in control and EpCAM-mutated CTE patients. D1-2, daughter crypts. Scale, bar, 50μm. **G**, Quantification of symmetric versus asymmetric crypt fissions in control and EpCAM-mutated CTE patients. Mean symmetric fission in controls = 90.14±15.29 % (mean±S.D), mean asymmetric fission in controls = 9.87±15.29 %, mean symmetric fission in CTE patients = 52.26±29.84 %, mean asymmetric fission in CTE patients = 47.74±29.84 %. n (control) = 12 patients, n = (CTE) = 13 patients. Two-way ANOVA, Tukey’s comparison test, *** p = 0.0001. **H-I**, Representative time-lapse sequences of crypt fission in control-KO (**H**) and EpCAM-KO (**I**) organoids. D1-2, daughter crypts. Scale, bar, 50μm. **J**, Statistical analysis of the number of crypt fission events per crypt in control-KO and EpCAM-KO organoids. Mean number of crypt fission per crypt in control-KO = 0.62±0.1 (mean±S.E.M), in EpCAM-KO = 1.42±0.26. Unpaired t test, ** p = 0.003. **K**, Statistical analysis of the percentage of symmetric or asymmetric crypt fissions in control and EpCAM-KO organoids. Mean percentage of symmetric crypt fission in control-KO = 33.63±15.22% (mean±S.E.M), in EpCAM-KO = 21.58±11.45%, mean percentage of asymmetric crypt fission in control-KO =66.37±15.22%, in EpCAM-KO = 78.41±11.45%. Unpaired t test, * p = 0.02, ** p= 0.001.

Analyzing in more detail the histological samples from *EPCAM*-mutated CTE patients, we noted a significant increase in the frequency of characteristic signs of crypt fission events compared to controls, rising from 7.16±1.65% (mean±S.E.M) in controls to 14.31±1.50% in CTE patients (Figure 6C-D). In addition, in control biopsies, crypt fission was predominantly symmetric, characterized by two future daughter crypts of similar size and shape, and with comparable surface lumina (Figure 6E-G). In contrast, CTE samples showed clear signs of asymmetric crypt fission, reflected by a significant shift in the ratio of symmetric to asymmetric fission events (Figure 6C blue arrowheads, E-G) and a frequent occurrence of multifid crypts (Figure 6C yellow arrowheads). Thus, the crypt fission process was altered under EpCAM’s loss in CTE patients. However, crypt fission frequency is known to be influenced by factors such age and physiological condition ^35,39,40,41,42^. Therefore, we could not exclude the possibility that crypt abnormalities observed in CTE biopsies resulted from patient-specific effects or environmental influences rather than directly from EpCAM loss in crypt epithelial cells. To definitely establish a direct role of EpCAM in crypt bifurcation, we tackled this process *in vitro* using control and EpCAM-KO 3D organoids. Live imaging of crypt remodeling was conducted for 72 h. Control crypt fission was completed within 40 h in average (Figure 6H; Video 8). In contrast, EpCAM loss caused an increase in crypt fission events, with the mean rate rising from 0.62±0.1 (mean±S.E.M) in controls to 1.42±0.26 in EpCAM-KO organoids (Figure 6J). Additionally, the proportionality of asymmetric crypt fission was more elevated in EpCAM-KO organoids compared to controls (Figure 6I,K; Video 9). These *in vitro* findings aligned with observations from patient biopsies (Figure 6C-G). Collectively, these findings demonstrated a critical role for EpCAM in regulating crypt fission both *in vitro* and *in vivo*, maintaining orderly progression through the crypt cycle and thereby ensuring the proper formation of two daughter crypts with identical morphology.

## DISCUSSION

In this study, we identified EpCAM as a key regulator of the mechanical homeostasis of the intestinal crypt. While EpCAM has been extensively studied for its functions in epithelial adhesion, signaling, and barrier integrity ^43,44,12,13^, its contribution to tissue mechanics within the intestinal stem cell niche has remained largely unexplored. Our work revealed that EpCAM is not a passive epithelial marker but instead acts as a central organizer of crypt mechanical properties, with particularly profound consequences for Paneth cell behavior and the fidelity of the crypt fission process (Figure 7).

**Figure 7:**
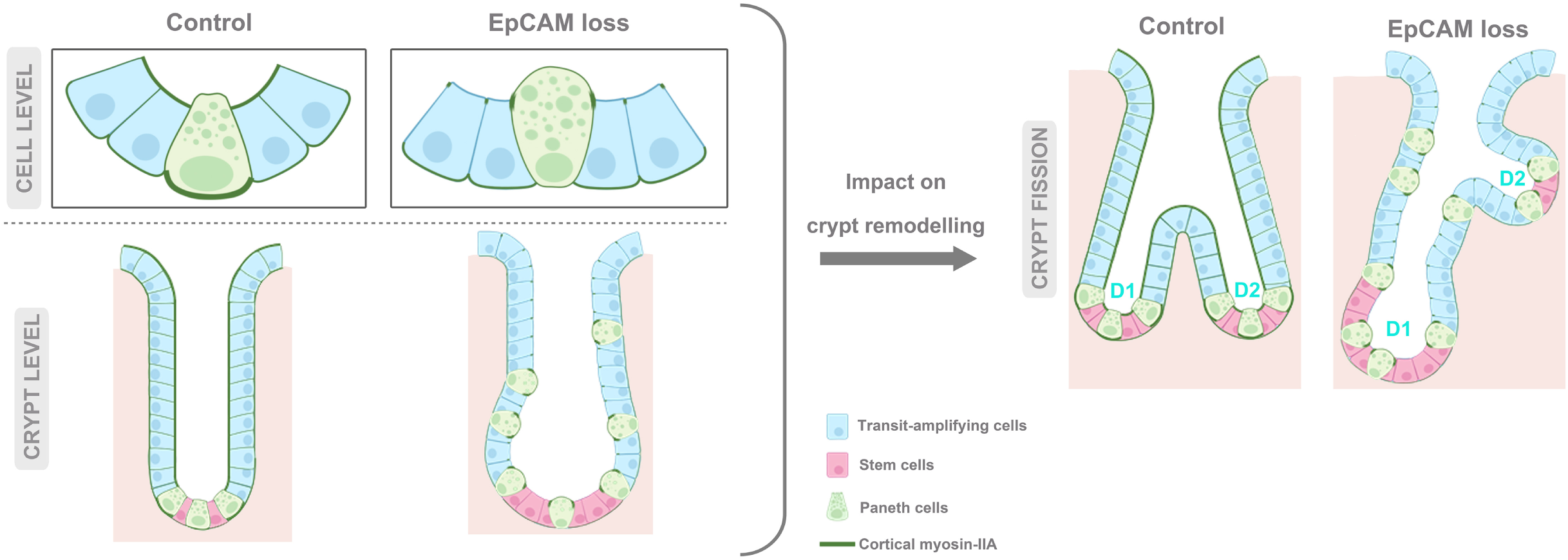
Scheme depicting the impact of EpCAM’s loss on crypt cell behavior and consequences on crypt organization and fission. D1-D2, daughter crypts.

### EpCAM acts as a regulator of crypt mechanical integrity

EpCAM has been implicated in tissue morphogenesis in various models. In the zebrafish injected with a morpholino directed against EpCAM, epithelial cells in the enveloping layer show slower epiboly movements ^45^. Moreover, in Xenopus, loss of EpCAM leads to epidermal disorganization and the formation of cell cluster lesions on the tissue surface during embryogenesis ^11^. In the intestine, the clinical significance of EpCAM is highlighted by the rare childhood enteropathy CTE, for which patients present characteristic mechanical lesions of the villi ^17,18,12^. In mouse, the EpCAM-KO model reproduces these villi architectural and functional defects ^46^. Surprisingly, no major tissue defects were reported in the intestinal crypts of either CTE patients or the EpCAM-KO murine model. In the mouse model, this can be explained by the fact that EpCAM-mutant mice show severe intestinal failure and die at 3-4 days after birth ^46^. At this postnatal stage, intestinal crypts are not yet present in mice and are not definitively formed until later, 7 days after birth ^25^. Whether CTE patients also display significant crypt abnormalities has not yet been directly investigated.

Here, we demonstrated that EpCAM regulates crypt biomechanics by modulating cellular forces at the crypt base. Functional perturbation of EpCAM disrupted global crypt rigidity and profoundly redistributed actomyosin contractility. AFM measurements revealed that EpCAM loss led to a significant modification in overall crypt stiffness (Supplementary Figure 1D-G). Although seemingly counterintuitive given EpCAM’s classical association with epithelial cell adhesion, EpCAM does not function as a canonical cadherin. Instead, it indirectly modulates junctional architecture through regulation of the actin cytoskeleton and actomyosin activity ^14,15,11,12,13^. Accordingly, the increased rigidity observed upon EpCAM silencing likely reflects misregulated cortical tension rather than strengthened cell–cell adhesion.

Consistent with this interpretation, EpCAM depletion drastically remodeled the crypt contractile network. Both apico-basal cortical pools and lateral localization of myosin II-A were disrupted, indicating a loss spatial organization of contractile forces within crypts (Figure 1F-G). Laser ablation experiments provided critical mechanistic insights, revealing that lateral cell–cell contacts at crypt base generate tensile forces, evidenced by junctional constriction following ablation (Figure 1K-L, N-O; Videos 3-4), but of a different nature than positive tension typically reported in other epithelial systems ^47,48,29^. This distinctive mechanical regime may reflect adaptations of columnar crypt epithelial cells to their tubular geometry and niche function, enabling cell elongation, dense epithelial packing, and resistance to luminal and extracellular mechanical stresses. Importantly, EpCAM depletion markedly attenuates the amplitude of these lateral tension forces (Figure 1N-O; Videos 6-7), demonstrating that EpCAM is required to sustain interfacial tension at crypt lateral contacts. This phenomenon is likely attributable to the perturbation of the mechanical interfacial tension within the crypt base, illustrated by the EpCAM-induced shift in the apico-basal gradient of myosin-IIA that establishes at lateral contacts in contrast with homogenous contractile distribution at cell interfaces in controls (Figure 1F, I-J). This EpCAM-KO phenotype could reflect a reduced pool of mobilizable myosin-IIA following laser ablation of the lateral contact. Alternatively, it may arise from defective or delayed activation of myosin-IIA at the lateral cortex, potentially due to impaired upstream RhoA signalling. Indeed, we have previously demonstrated that EpCAM is required for the endosomal turnover of RhoA-GTP at the epithelial cortex ^13^. Together, these perturbations disrupted crypt mechanical stability and highlighted a unique biomechanical regulatory principle, whereby apical constriction, lateral contractility and interfacial tension must be tightly coordinated to preserve crypt geometry and organization. Our findings identified EpCAM as a central orchestrator, ensuring that cytoskeletal tension is properly oriented and balanced to support crypt architecture.

Tissue mechanics has emerged as a fundamental determinant of crypt morphogenesis, particularly during crypt budding and initial outgrowth ^25,24,7,49^. Despite its major impact on epithelial cell mechanics, EpCAM loss did not affect early stages of crypt formation, such as initial budding and elongation (Supplementary Figure 2B), but severely compromised crypt maturation and geometry at later stages (Figure 2A-D; Supplementary Figure 2B). Moreover, EpCAM-KO crypts became elongated, distorted, and irregular - a phenotype faithfully recapitulated in CTE patient biopsies (Figure 2H-O). This temporal uncoupling suggests that mechanical requirements may evolve during crypt development. Recent work has emphasized that intestinal morphogenesis relies on dynamic coupling between proliferation, differentiation, and mechanical forces ^50,24^. Our findings place EpCAM at the core of this coupling, ensuring mechanical conditions conductive to straight crypt elongation and architectural stability. EpCAM-based mechanical coordination sustained both crypt shape, and the timing and symmetry of crypt fission. Importantly, despite the pronounced morphological defects, crypt–villus compartmentalization, differentiation, and stem cell niche organization are largely preserved (Supplementary Figures 3-4). Moreover, OLFM4^+^-stem cells remain properly positioned, and proliferative compartments are maintained, indicating that key biochemical signaling pathways (e.g., Wnt, Notch, BMP) are largely intact, and that the observed phenotype upon EpCAM loss arises primarily from mechanical dysregulation rather than altered cell fate specification. This distinction is critical, as prevailing models of crypt homeostasis largely emphasize biochemical signaling gradients and stem cell dynamics. Our findings instead support a model in which correct crypt architecture emerges from the integration of biochemical cues with finely tuned mechanical programs, and in which proper crypt maturation requires not only molecular patterning but also precise mechanical homeostasis, which is orchestrated in part by EpCAM.

### Paneth cells are mechanically specialized organizers of the crypt niche

At the level of niche organization, Paneth cells serve as both biochemical and physical organizers of the crypt base ^31,32,51^. In our study, Paneth cells also emerge also as critical mediators of EpCAM-dependent mechanics. Paneth cells exhibit higher levels of cortical myosin-IIA and more pronounced contractile asymmetry compared to other crypt cells (Figure 3F-H; Supplementary Figure 5A). EpCAM is enriched at Paneth cell interfaces (Figure 3A-E), and its loss selectively disrupts Paneth cell mechanical homeostasis, leading to altered cortical myosin-IIA distribution (Figure 3F-K). At the cellular level, EpCAM’s silencing led to an impairment of canonical pyramidal morphology of Paneth cells, their apical domain expansion (Figure 4A-D), and pronounced changes in their cell aspect ratio (Figure 4D-I) and coordinated alignment with other crypt cells within the crypt base (Figure 4A-B, 4H-L). Collectively, these cell defects may explain why Paneth cells lose their tight clustering in crypt bases, and why some were found far from the crypt bottom in EpCAM mutants (Figure 5; Supplementary Figure 7). Alternatively, crypt dilatation observed under EpCAM mutation may also participate in the scattering of Paneth cells in crypts (Figure 1F-G; Figure 2H-J,M; Figure 5; Supplementary Figure 7). Indeed, a recent study from K. Anseth and colleagues demonstrated that the density of Paneth cells relies on the degree of crypt curvature, where highly curved crypts concentrate Paneth cells at their base ^52^.

Based on our data, we propose that EpCAM-dependent mechanical homeostasis of Paneth cells may be required for their function as structural organizer of the crypt base, and that its disruption may underlie the crypt architectural defects observed in EpCAM-deficient tissues. Altered Paneth cell morphology increased variability in cellular orientation and disrupted collective epithelial order in the crypt (Figure 4H-L). These cellular defects translated to tissue-scale consequences, including irregular crypt geometry and overall crypt disorder, highlighting Paneth cells as potential mechanical hubs that integrate single-cell contractility with tissue-level architecture. Our data therefore support a model in which Paneth cells act not only as biochemical niche cells, but also as mechanical organizers that contribute to shaping crypt architecture through force generation and spatial organization.

### EpCAM-dependent mechanics as a driver of crypt fission fidelity

Beyond maintaining steady-state architecture, EpCAM-mediated Paneth cell mechanics play a critical role in the morphogenetic process of crypt fission. Crypt fission is a fundamental morphogenetic process that drives intestinal expansion - during embryogenesis in humans and postnatal maturation in mice ^53,54,35^ – and, to a lesser extent, contributes to tissue homeostasis in adulthood ^55,56^. Traditionally, the accepted crypt cycle model describes crypt fission as a slow, continuous process initiated in the stem cell compartment at the crypt base ^53,56^. First, the crypt grows in size to an equivalent to twice the initial pool of stem cells compacted at the crypt bottom ^34,35^. Subsequently, mechanical buckling of the epithelial layer at the crypt base generates tissue buds that evolve into a symmetrical bifurcation, progressing upward until full fission produces two clonal crypts ^57,34^. Crypt fission may be influenced by cell division orientation: Quyn et al. showed that *APC* mutation disrupts the preferential orientation of cell divisions, leading to aberrant cell shapes and defective crypt architecture ^58^. Extending this, Boman and Fields proposed that loss of spindle orientation in Apc-deficient contexts drives uncontrolled expansion of the crypt base, leading to increased asymmetric fission and formation of disorganized crypts and microadenomas ^37^.

Alternative models emphasize mechanics more directly. For instance, computational simulations by Pin et al., suggested that crypt fission could be initiated by different mechanical properties between Paneth and stem cells, independently of proliferative growth ^59^. Additionally, Näthke and collaborators have reported that proper positioning and ratios of stem cells and Paneth cells at the crypt base are required for correct crypt fission ^33^. Paneth cells may mechanically constrain neighboring stem cell clusters, creating local regions of increased deformability and epithelial buckling ^33^. Moreover, Paneth cells could provide mechanical resistance within the crypt base ^33^, although this has not been demonstrated experimentally. Here, our data demonstrated that EpCAM-dependent contractility of Paneth cells is a central determinant of the crypt fission process. In both CTE patient biopsies and EpCAM-KO organoids, crypt fission events were more frequent, with a random organization, indicating disrupted bifurcation dynamics (Figure 6). Live imaging in organoids confirmed that EpCAM deficiency increases crypt fission rates and the probability of asymmetric daughter crypt formation (Figure 6H-K; Video 9), directly linking molecular regulation of contractility to crypt-scale remodeling. These findings demonstrated that the mechanical integrity of Paneth cells, regulated by EpCAM, acts as a key driver of crypt bifurcation and ensures symmetric propagation of the crypt network.

### Implications for epithelial organization and disease

The profound impact of EpCAM loss on crypt fission dynamics may have broader pathological implications. Although physiological crypt fission is underexplored, it is known to be engaged during epithelial regeneration following cytotoxic injury ^39^, irradiation ^40,41^, or surgical resection^42^. Pathological disruptions in the crypt cycle can lead to an asymmetrical outcome and corrupted crypts, a phenomenon linked to adenocarcinoma development in the small intestine and colon ^60,61,57^. This link is well established in cases of *APC* mutations, as seen in Familial Adenomatous Polyposis (FAP) patients and *APC*^⁺/⁻^ mice, where crypt disorganization is observed ^38,62^. EpCAM’s role in tumorigenesis is debated ^63^. Originally identified as a tumor marker, it is aberrantly expressed in various carcinomas (e.g., colorectal, breast) and used as a diagnostic marker ^64^. Clinical data also associate EpCAM with epithelial dysplasia and tumor development. In Lynch syndrome, *EPCAM* deletions disrupt its 3’ end, leading to transcriptional read-through and inactivation of the adjacent MSH2 gene—a DNA repair factor—thereby predisposing to colorectal cancer ^65,66,67^. The studies performed so far only concern the consequences of *MSH2* inactivation, and do not take in account the mechanical consequences directly linked to the disruption of EpCAM expression. Interestingly, in the pancreatic epithelial tissue, an imbalance of the apico-basal tension gradient has also been observed and directly correlated with tumor drift ^2^. Thus, the loss of mechanical homeostasis at cellular interfaces in crypt bases following EpCAM loss, together with the resulting abnormalities in crypt fission geometry, could at least partly account for the role of EpCAM in protecting from intestinal pre-tumoral drift.

## Acknowledgments

We thank Guillaume Salbreux and Quentin Vagne (UNIGE, Geneva), and the Cell Adhesion and Mechanics team (IJM) for helpful discussions. We thank Jad Saleh (Institut Jacques Monod, IJM) and Pierre Nouaux (Institut de Biologie du Développement de Marseille, IBDM) for their technical help. Confocal microscopy analyses were performed in the ImagoSeine microscopy facility (IJM) and PICSL imaging facility (IBDM). This work was supported by grants from the Groupama Foundation – Research Prize for Rare Diseases 2017 (to D.D), the Human Frontier Science Program (RGP0038/2018) (to D.D.), the CNRS through the MiTi interdisciplinary programs (to D.D.), the Ligue contre le Cancer (to L.R.), the European Commission (MSCA to A.B.), the Fondation ARC (to M.S., F.R. and D.D), the Agence Nationale de la Recherche (ANR-25-CE13-3505-01 to D.D.), and the 2021-2023 Cancer Control strategy, on funds administered by INSERM (to D.B. and D.D.).

## Author contributions

M.S., L.R., D.B., A.B., B.L., S.R., V.O., T.D., and D.D. designed and performed experiments and required analyses. M.S., L.R., D.B., J.S., F.R., F.R. and D.D. coordinated the overall research and experiments, and wrote the manuscript.

## METHODS

Animal procedures were conducted in accordance with the guidelines of the French regulation for animal care. No ethical approval was required, as animal procedures were restricted to mouse sacrifice for organ dissection. Human tissue samples were obtained according to a protocol approved by the Comité de Protection des Personnes (CPP) (#2014-01-04MS6). All parents signed informed consent forms approved by the local ethics committee for biopsy exploitation (Unité de Recherche Clinique (URC) of Necker Hospital, URC).

### Mouse models

Wild-type C57/Bl6 adult male mice were provided by the animal house facility of the Institut de Biologie du Développement de Marseille (Marseille). VillinCreERT2-tdTomato organoids were generated from mice provided by Danijela Vignjevic (Insitut Curite, Paris) ^68,69^. Myosin-IIA-KI-GFP organoids were generated from myosin-IIA-GFP-knock-in mice provided by Robert S. Adelstein (NHLBI, Bethesda) and Ana-Maria Lennon-Dumesnil (Institut Curie, Paris) ^57,69^. Mice were housed in EOPS (Environment without Specific Pathogenic Organisms) environment, and handled in accordance with French regulation for animal care.

### Organoid cultures and transfection

#### 3D organoid cultures

3D organoid cultures were generated and maintained as previously described ^69^. Small intestinal crypts were isolated from 6- to 12-week-old adult male mice. After euthanasia by cervical dislocation, the small intestine was harvested, flushed with cold PBS to remove luminal content, opened longitudinally and cut into 3-5 mm pieces. Tissue fragments were washed thoroughly in cold PBS, incubated on ice in 5 mM EDTA for 10 min, vortexed for 2 min to release villi, and the supernatant was then collected as ‘Fraction 1’. After EDTA removal, the intestinal pieces were vigorously vortexed for 3 min in cold PBS to release remaining villi and crypts (‘Fraction 2’). This process was repeated 2 times (‘Fractions 3 and 4’). The fractions 3-4 were concentrated in crypts, so these were pooled, filtered through a 70-µm cell strainer to remove tissue debris and centrifuged at 1000 rpm for 5 min. Pelleted crypts were washed in advanced DMEM/F12 (#12634010 Thermo Fisher Scientific) and centrifuged again. The final pellet was resuspended in ice-cold Matrigel (#734-1100 VWR) and plated as domes. Incubation at 37°C for 20-30 min allowed Matrigel polymerization.

3D organoids were cultured in IntestiCult™ ENR Organoid Growth medium (#06005 StemCell Technologies). Organoids were passaged every 5 to 6 days by mechanical dissociation, and culture medium was refreshed every 2 days. Live-imaging or immunofluorescence experiments were performed from day 3 to 5 post-seeding.

#### Organoid-derived monolayer cultures

Organoid-derived monolayers were generated as previously described ^30,70^. L-WRN conditioned medium was prepared as previously described by Stappenbeck and colleagues ^71^, using L-WRN cells obtained from ATCC (#CRL-3276). L-WRN cells were cultured in L-cell medium, consisting of high-glucose DMEM (Sigma-Aldrich, #D6429) supplemented with 10% fetal bovine serum (FBS, #A5256701 Gibco), 2 mM GlutaMAX (#35050-038 Gibco), and 100 U/mL penicillin-streptomycin (#10378016 Gibco). After 24 hours, selection antibiotics were added: 500 μg/mL geneticin (#10131-027 Gibco) and 500 μg/mL hygromycin (#10687010 Gibco). Once confluent, cells were passage into five T175cm^2^ flasks and expanded in L-cell medium until they reached confluence again. At confluence, cells were switched to Primary Cell Medium (PCM), composed of Advanced DMEM/F12 supplemented with 20% FBS, 2 mM GlutaMAX, and 100 U/mL penicillin-streptomycin. Conditioned PCM — enriched with secreted Wnt3a, R-spondin, and Noggin — was collected every 24 hours over several days. Each collected batch was mixed 1:1 with freshly prepared PCM, then vacuum-filtered through a 0.22 μm membrane filter to generate the final L-WRN conditioned medium. 3D organoids were cultured in L-WRN conditioned media for at least 3 days before use for the generation of organoid-derived monolayers.

To produce cross-linked Matrigel (CL-Matrigel) substrates, a fresh-made cross-linker solution was prepared by mixing 100 mM NHS (#130672Sigma-Aldrich) and 400 mM EDC (#E1769 Sigma-Aldrich) in cold PBS 4°C. Glass coverslips (Ø = 18 mm) were plasma-treated and cooled at -20°C for 3 min. Cross-linker solution was mixed with pure Matrigel at a ratio of 1:10 (v/v) and 50µL drops of the mixture were poured on top of cooled plasma treated coverslips and spread by tilting the coverslips. Then, coverslips were placed in a 37°C incubator in a 12 well plate for 2h to form CL-Matrigel layers. PBS was poured in 2 empty wells of the 12 well plate ware to prevent gel dehydration. The CL-Matrigel substrates were washed with PBS and incubated in PBS at 37°C for 24 h to remove unreacted EDC and NHS. CL-Matrigel substrates can be stored up to a week in PBS at 37°C for future use.

Before use, CL-Matrigel coated coverslips were washed twice in PBS. 3D organoids were harvested with cold advanced DMEM/F12. Organoids were mechanically broken through a P200-filtered tip 150 times and centrifuged at 72g for 3 min at 4°C. The supernatant was removed and 5 mL of fresh F12 was added to the pellet. Breaking and centrifugation steps were repeated once. Then, cell pellet was filtered through a 30 µm cell strainer and resuspended in warm L-WRN conditioned medium complemented with 10µM Y27632 (StemCell #72302) and 150 µL were gently seeded on the CL-Matrigel substrate. After 4h incubation at 37°C, up to 1 mL/well of the L-WRN conditioned medium / Y27632 solution was added for the first 24h culture. After 24h, cells were grown on L-WRN conditioned medium (without Y27632). Culture media were refreshed every 24-48 hours with L-WRN conditioned medium. Organoid-derived monolayers were cultured for 10 days.

### Organoid transfection

EpCAM-KO organoids were generated by a CRISPR-Cas9 strategy using the Edit-R lentiviral sgRNA system from Horizon Discovery (Cambridge, UK). A set of 3 lentiviral sgRNAs were used to target *EPCAM* (#GSGM11839-246765098: target sequence TAGTAAATCAGAGTCTGCCC; #GSGM11839-246765101: target sequence CTTGTCGGTTCTTCGGACTC; #GSGM11839-246765094: target sequence GGGCGATCCAGAACAACGAT). Control-KO organoids were generated using a non-targeting sgRNA (#GSG11811, Horizon Discovery).

Organoid-derived monolayers were first transduced with lentiviral particles encoding a doxycycline-inducible Cas9 construct. After selection with 2 μg/ml of blasticidin (#BP2647, Fisher Scientific) for three days, a second transduction was performed using lentiviruses encoding the sgRNAs. A subsequent round of selection was carried out with 2 μg/ml of puromycin (#A1113803, Thermo Fisher Scientific) for three days. Cas9 expression was induced by addition of 500 ng/mL doxycycline (#D9891, Merck) to the culture medium. EpCAM deletion was confirmed by western blot and immunofluorescence.

### Patient biopsies and preparation of tissue samples

Tissue samples of *EPCAM*-mutated CTE children were provided by Necker-Enfants Malades, Robert Debré and Trousseau hospitals (Paris, France), and were collected either from the Necker Paediatric Anatomo-Pathology Department for retrospective analyses, or from the Gastroenterological functional exploration Departments for prospective studies. Genetic characterization of the CTE patients included here had been conducted in a previous study ^18^. The patient cohort analyzed here comprised *N* = 13 CTE patients mutated for *EPCAM* and aged 4 months to 17 years, as well as *N* = 12 controls (non CTE patients: N = 2 patients with Crohn’s disease, *N* = 1 patient with hyperinsulinemia, *N* = 1 patient with duodenal atresia, *N* = 2 patients with auto-immune enteropathies, *N* = 2 patients with gastro-esophageal reflux, *N* = 4 patients with coeliac disease) aged 9 months to 14 years, who underwent duodenal endoscopy for routine explorations. The CTE patients displayed typical CTE epithelial abnormalities in intestinal biopsies and parenteral nutrition dependency, and presented *EPCAM* loss-of-function mutations. Duodenal biopsies were collected during endoscopic procedures for diagnosis and/or monitoring of CTE and control infants. Human tissue samples were obtained according to a protocol approved by the Comité de Protection des Personnes (CPP) (#2014-01-04MS6). All parents signed informed consent forms approved by the local ethics committee for biopsy exploitation (Unité de Recherche Clinique (URC) of Necker Hospital, URC).

### Antibodies and reagents

Mouse monoclonal (#TA506627, IF dilution 1:100) antibody directed against human EpCAM was from Origene (Rockville, USA). Mouse monoclonal antibody directed against CK-20 (#M7019, IF dilution 1:100) was from Dako. Rat monoclonal antibody directed against mouse EpCAM (#G8.8, IF dilution 1:20) was from DSHB. Rat monoclonal antibody directed against E-cadherin (#U3254, IF dilution 1:100) was from Sigma-Aldrich. Rabbit polyclonal antibodies directed against lysozyme were from Dako (#A0099, IF dilution 1:200) and Abcam (#ab108508, IF dilution 1:200). Rabbit polyclonal antibodies directed against myosin-IIA (#3403S, IF dilution 1:100), against OLFM4 (#39141S, IF dilution 1:100), or against Ki67 (#9129, IF dilution 1:100), were from Cell Signaling. Rabbit polyclonal antibody directed against EpCAM (#ab71916, IF dilution, 1:100) was from Abcam. Goat anti-mouse-Alexa-488, -568, anti-rabbit-Alexa-488, -568 or -647, anti-rat-Alexa-488 were from Invitrogen (Paisley, UK). Nuclei were stained with Hoechst 33342 solution (Life Technologies) at a 1:1000 dilution.

### Biochemical analysis

For western blots, organoid lysates were prepared 3 days after plating using for each condition 6 wells of a 24-well plate. Matrigel was depolymerized by incubating with 1 ml of Gentle Cell Dissociation Reagent (#07174 StemCell Technologies, Vancouver, Canada) for 30 min at 4°C and centrifugation for 5 min at 500 x g at 4°C. The pellet was resuspended in lysis buffer containing 25mM Tris / 5 mM NaCl / 1mM EDTA / 1mM EGTA / 0.5% NP-40 / 1% Triton TX100, and incubated on ice for 30 min. The solution was then passed 10 times through a syringe equipped with a 23G needle and centrifuged at 10 000 RPM at 4°C for 10 min. Supernatant total protein content was measured by Bradford assay (#5000205 Biorad). For each condition, 50μg of proteins was loaded per well in Novex Tris-Glycine pre-cast gels (ThermoFischer Scientific). Proteins were detected with rabbit polyclonal antibody directed against EpCAM (Abcam #ab71916, dilution 1:500) or mouse monoclonal antibody directed alpha-tubulin (Sigma-Aldrich #T6199, dilution 1:500), and HRP-linked donkey anti-rabbit IgG antibody (Invitrogen #A16035, dilution 1:10,000) or HRP-linked goat anti-mouse IgG antibody (Sigma-Aldrich #12-349, dilution 1:10,000), respectively, and visualized on ImageQuant LAS4000 (GE-Healthcare, Buckinghamshire, UK). Signal quantification was performed using Fiji software.

### Immunostaining

Routinely, 3D organoids and organoid-derived monolayers were fixed using 4% paraformaldehyde for 30 min, then permeabilized using 0.025% saponin solution in PBS for 30 min. Blocking step was performed in 0.025% saponin/1% BSA solution for 45 min, before proceeding to incubation with primary antibody at 4°C overnight. The next day, the primary antibody was removed and the organoids washed 3 times in PBS for 10 min each, before adding the secondary antibody and left to incubate for 2h at room temperature. Finally, organoids were washed 3 times again for 10 minutes before incubating in Hoechst 33342 for 15 min to stain nuclei. Immunostained samples were mounted in home-made Mowiol solution. For organoid proliferation analysis, 5ug EdU (Thermo Fischer scientific, #A10044) was delivered in the culture medium for 15min prior to PFA cell fixation. To label proliferative cells, organoid monolayers were stained with the Click-iT EdU Alexa Fluor 555 Imaging kit (Thermo Fischer scientific, #C10338).

For fluorescent immunostaining *in vivo*, mouse jejunum or human duodenum biopsies were processed as previously described ^34;58^. Briefly, samples were fixed for 2 hours in 4% PFA and paraffin embedded. 5 μm tissue sections were de-waxed in a xylene bath, rehydrated in isopropanol and in solutions with decreasing ethanol concentrations. Tissue sections were then blocked in 10% goat serum (#S26-M Sigma-Aldrich) for 1 h. Primary antibody incubation was performed at 4°C overnight and secondary antibody incubation at room temperature for 2 h, both in 1% goat serum solution. Hoechst33342 staining was used to detect nuclei. Tissue sections were mounted in home-made Mowiol-488 solution.

Immunohistological stainings were performed as described in ^59^. Briefly, 5μm paraffin sections were treated with 3% hydrogen peroxide (#H1009, Sigma-Aldrich) and antigen retrieval was performed in citrate buffer pH6 (#H-3300, VectorLab). Tissue sections were then incubated with the avidin/biotin blocking ABC kit (#PK-6100, VectorLab), and immunoreactions were revealed with DAB substrate (#SK-4100, VectorLab).

### Stiffness measurements of live organoid crypts

4-day post-seeding organoids were used for AFM-based stiffness measurements. Organoids were gently released from Matrigel domes and laid on a thin polymerized layer of Matrigel with the crypt surface left exposed. After letting them attach for at least 30 min, culture medium was added, and organoids were allowed to stabilize for 1 hour in a standard incubator prior to measurement. AFM measurements were performed using Nanowizard-IV system (JPK/Bruker, Germany) mounted on an inverted optical microscope (Nikon, Japan). A soft cantilever probe (SAA-SPH-1UM, Bruker) with a nominal spring constant of ∼0.25 N/m with a cylindrical tip of 1 μm radius was used. The AFM tip was positioned above the exposed crypt region, and force–distance indentation curves were acquired. The Young’s modulus (*E*) was estimated by fitting the Hertz model to the approaching trace of the resulting force-indentation (*F-δ*) curves using JPK SPM data processing software (Bruker), assuming a spherical indenter geometry of radius (*R*) and Poisson’s ratio (*ν*) of 0.5 ^72^ :

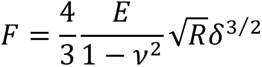

### Organoid live imaging and laser ablation

Live imaging of intestinal organoids was performed to monitor crypt fission dynamics over time. Imaging was carried out using an AxioObserver Z1 Colibri DEF2 inverted microscope (Zeiss), equipped with a 10x dry objective, under standard culture conditions (37 °C, 5% CO₂) for up to 60 hours. Organoids were imaged in time-lapse mode with 15-min intervals, and maintained in a sealed stage-top incubator throughout the acquisition to ensure stable environmental control.

For laser ablation experiments, membranes were first labelled in live organoid-derived monolayers by incubating with CellMask™ Orange Membrane Tracking stain (Invitrogen #C10045) for 3 h in culture media. Lase ablations on labelled organoids were performed using a Zeiss LSM 880 microscope equipped with a tunable two-photon laser set to 780 nm at 20% power for 5 iterations, based on ^73^. Lateral membrane lengths were measured before and after laser ablation using ImageJ.

### Segmentation, 3D rendering and analyses

Three-dimensional cell segmentation was performed on confocal z-stacks using Cellpose algorithm (v3) (https://www.cellpose.org/) in 3D mode with the Zstack option enabled. Segmented volumes were loaded into Napari (v0.4.18) (https://napri.org) for quality control and manual correction of individual masks when necessary. Each cell label within the 3D volume was then extracted as an independent binary mask using a custom Python script. For each mask, voxel dimensions were preserved to ensure spatial accuracy. Binary masks were subsequently converted into 3D surface meshes using a Python code (available at Zenodo: 10.5281/zenodo.7802531), as previously described ^69^. Final 3D renderings and mesh visualizations were performed in MeshLab (https://www.meshlab.net), allowing for structural inspection, surface smoothing, and figure generation. This pipeline enabled high-resolution reconstruction of individual epithelial cells within crypt-like domains for shape analysis.

To assess nucleus orientation, three-dimensional image stacks were analyzed using manually segmented nuclear masks together with a reference structural signal defining the tissue axis. Two-dimensional projections were generated to obtain a continuous representation of this structure, from which the external contour was extracted and smoothed.

For each nucleus, the principal axis of elongation was determined by principal component analysis (PCA) applied to its three-dimensional voxel coordinates. Nuclear centroids were projected onto the imaging plane. A local reference direction was defined as the tangent to the tissue contour at the closest point to each nucleus, refined based on local signal intensity to improve robustness. This tangent was consistently oriented using a predefined base point to ensure uniform polarity across the dataset. To resolve the directional ambiguity of PCA, nuclear axes were oriented relative to a user-defined reference line. The angle between the oriented nuclear axis and the local tangent was then calculated in the range of 0° to 180°. In cases where both directions were nearly aligned, a correction step was applied to stabilize axis orientation. To assess local coordination, a nearest-neighbor analysis was performed in three dimensions using nuclear centroids. Neighboring cells were defined within a fixed distance threshold (110 pixels), which was empirically determined by comparing the number of neighbors observed manually in the images with those identified computationally, and adjusting the parameter until both approaches were consistent. Alignment between neighboring nuclei was quantified using the absolute dot product of their principal axes. All analyses were performed using custom Python scripts based on standard scientific computing libraries.

### Statistical analysis

All statistical analyses were performed using Prism v10 (GraphPad Software, San Diego, CA, USA). The specific statistical tests used, sample sizes, and p-values are indicated in the corresponding figure legends. Unless otherwise specified, all experiments were independently replicated at least three times.

### Creation of graphics

Graphics have been generated with BioRender software.

### Reporting summary

Further information on research design is available in the Nature Portfolio Reporting Summary linked to this article.

## Data availability

Data generated in this study are provided in the Source Data. Microscopy data, and Python scripts for nuclei vector orientation reported in this paper will be shared by the lead contact on request. Any additional information required to reanalyze the data reported in this paper is available upon request from the lead contact, Delphine Delacour.

## Materials availability

Materials generated in the current study are available from the lead contact upon request. There are restrictions to the availability of materials due to collaborations or MTAs.

## SUPPLEMENTARY INFORMATION

Supplemental Figures 1-7 and Videos 1-9 are available in the online version of the paper.

## VIDEOS

**Video 1: Behavior of a crypt cell after laser ablation in the cytoplasm**

**Description:** Time-lapse imaging of cell behavior after laser ablation within the cytoplasm in a crypt labelled CellMask-Membranes-GFP. Laser ablation is indicated with red circle at 5 sec. Scale bar 10 µm. Images were acquired every 1 s. Frame rate is 7 fps.

**Video 2: Behavior of a crypt cell after laser ablation of basal plasma membrane**

**Description:** Time-lapse imaging of cell behavior after laser ablation of the basal membrane in a crypt labelled CellMask-Membranes-GFP. Laser ablation is indicated with red circle at 5 sec. Scale bar 10 µm. Images were acquired every 1 s. Frame rate is 7 fps.

**Video 3: Behavior of a crypt cell after laser ablation of lateral plasma membrane**

**Description:** Time-lapse imaging of the behavior of a lateral contact after laser ablation in a crypt of CellMask-Membranes-GFP. Laser ablation is indicated with yellow circle at 5 sec. Scale bar 10 µm. Images were acquired every 1 s. Frame rate is 7 fps.

**Video 4: Behavior of a crypt cell before and after laser ablation of lateral plasma membrane**

**Description:** Time-lapse imaging of the behavior of a lateral contact after laser ablation in a crypt with CellMask-Membranes-GFP. Laser ablation is indicated with yellow circle at 5 sec. Membranes are shown in magenta before laser ablation, and in green after ablation. Scale bar 10 µm. Images were acquired every 1 s. Frame rate is 7 fps.

**Video 5: Behavior of myosin-IIA after laser ablation of lateral plasma membrane in an organoid crypt**

**Description:** Time-lapse imaging of the behavior of a lateral contact after laser ablation in a crypt of myosin-IIA-KI-GFP organoids. Laser ablation is indicated with yellow circle at 5 sec. Scale bar 10 µm. Images were acquired every 1 s. Frame rate is 7 fps.

**Video 6: Behavior of a crypt cell after laser ablation of lateral plasma membrane upon EpCAM-KO**

**Description:** Time-lapse imaging of the behavior of a lateral contact after laser ablation in a crypt of EpCAM-KO organoids stained with CellMask-Membranes-GFP. Laser ablation is indicated with yellow circle at 5 sec. Scale bar 10 µm. Images were acquired every 1 s. Frame rate is 7 fps.

**Video 7: Behavior of a crypt cell before and after laser ablation of lateral plasma membrane upon EpCAM-KO**

**Description:** Time-lapse imaging of the behavior of a lateral contact after laser ablation in a crypt of EpCAM-KO organoids stained with CellMask-Membranes-GFP. Membranes are shown in magenta before laser ablation, and in green after ablation. Laser ablation is indicated with yellow circle at 5 sec. Scale bar 10 µm. Images were acquired every 1 s. Frame rate is 7 fps.

**Video 8: Crypt fission behavior in control-KO organoids**

**Description:** Time-lapse imaging of crypt fission in control-KO organoids. Scale bar XXX µm. Images were acquired every 15 min. Frame rate is 40 fps.

**Video 9: Crypt fission behavior in EpCAM-KO organoids**

**Description:** Time-lapse imaging of crypt fission in EpCAM-KO organoids. Scale bar XXX µm. Images were acquired every 15 min. Frame rate is 40 fps.

## SUPPLEMENTARY INFORMATION

**Supplementary Figure 1:**
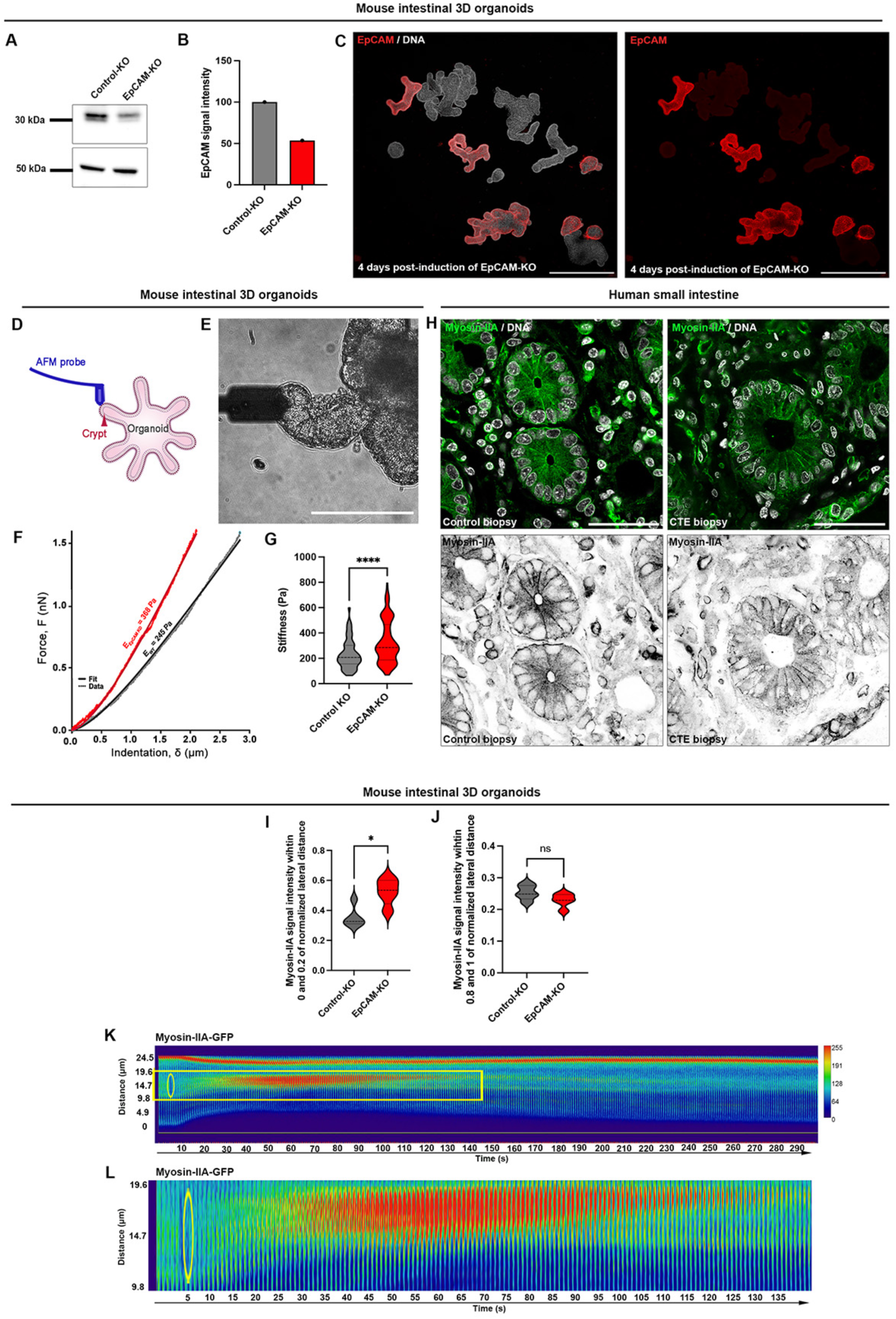
**A,** Western blot analysis of EpCAM expression in control-KO and EpCAM-KO organoids 3 days after induction. α-tubulin was used as a loading control. **B**, Analysis of EpCAM expression in control-KO and EpCAM-KO organoids. Mean EpCAM expression in EpCAM-KO organoids = 53.5%. **C**, Confocal microscopy analysis of EpCAM expression level in the EpCAM-KO organoid line 4 days after KO induction. Nuclei are stained with Hoechst 33342 (gray). Scale bar, 300μm. **D**, Schematic representation showing the principle of atomic force microscope (AFM) stiffness measurement. **E**, Phase contrast image showing AFM probe positioned on top of an organoid crypt (right). Scale bar, 100μm. **F,** Representative force-indentation curves for a control and an EpCAM-KO organoid crypt. **G,** Mean cortical stiffness of control and EpCAM-KO organoid crypts. Mean stiffness of control crypts = 233.2±10.48 Pa (mean± SEM), of EpCAM-KO crypts = 326.7±14.93 Pa. n (control-KO) = 121 crypts, n (EpCAM-KO) = 126 crypts. Mann-Whitney test, **** p < 0.001. **H**, Confocal analysis of myosin-IIA (green) distribution in crypts of control or EpCAM-mutated CTE patients. Nuclei are shown in grey. Scale bar, 50 μm**. I,** Statistical analysis of the signal intensity of myosin-IIA-GFP at the most apical part of the lateral contact (between 0 and 0.2 of normalized distance from the apical cell side). Mean intensity in control-KO = 0.35±0.03 (mean± SEM), in EpCAM-KO = 0.52±0.04. Mann-Whitney test, * p = 0.016. n (control-KO) = 24 contacts, n (EpCAM-KO) = 23 contacts. **J,** Statistical analysis of the signal intensity of myosin-IIA-GFP at the most basal part of the lateral contact (between 0.8 and 1 of normalized distance from the apical cell side). Mean intensity in control-KO = 0.25±0.01 (mean± SEM), in EpCAM-KO = 0.23±0.01. Welch’s test, ns non significative. n (control-KO) = 24 contacts, n (EpCAM-KO) = 23 contacts. **K,** Time-lapse images of myosin-IIA-KI-GFP control-KO organoids before or after laser ablation of a lateral contact. Laser ablation occurred at t = 5s. Yellow circles point toward the laser ablation site. Signal intensity is color-coded with Physics LUT table from ImageJ. Color scale bar indicates the gray value intensity. **L,** t-stack frame of the laser-ablated lateral contact was selected from a time-lapse series of myosin-IIA-KI-GFP organoids and presented as a kymograph.

**Supplementary Figure 2:**
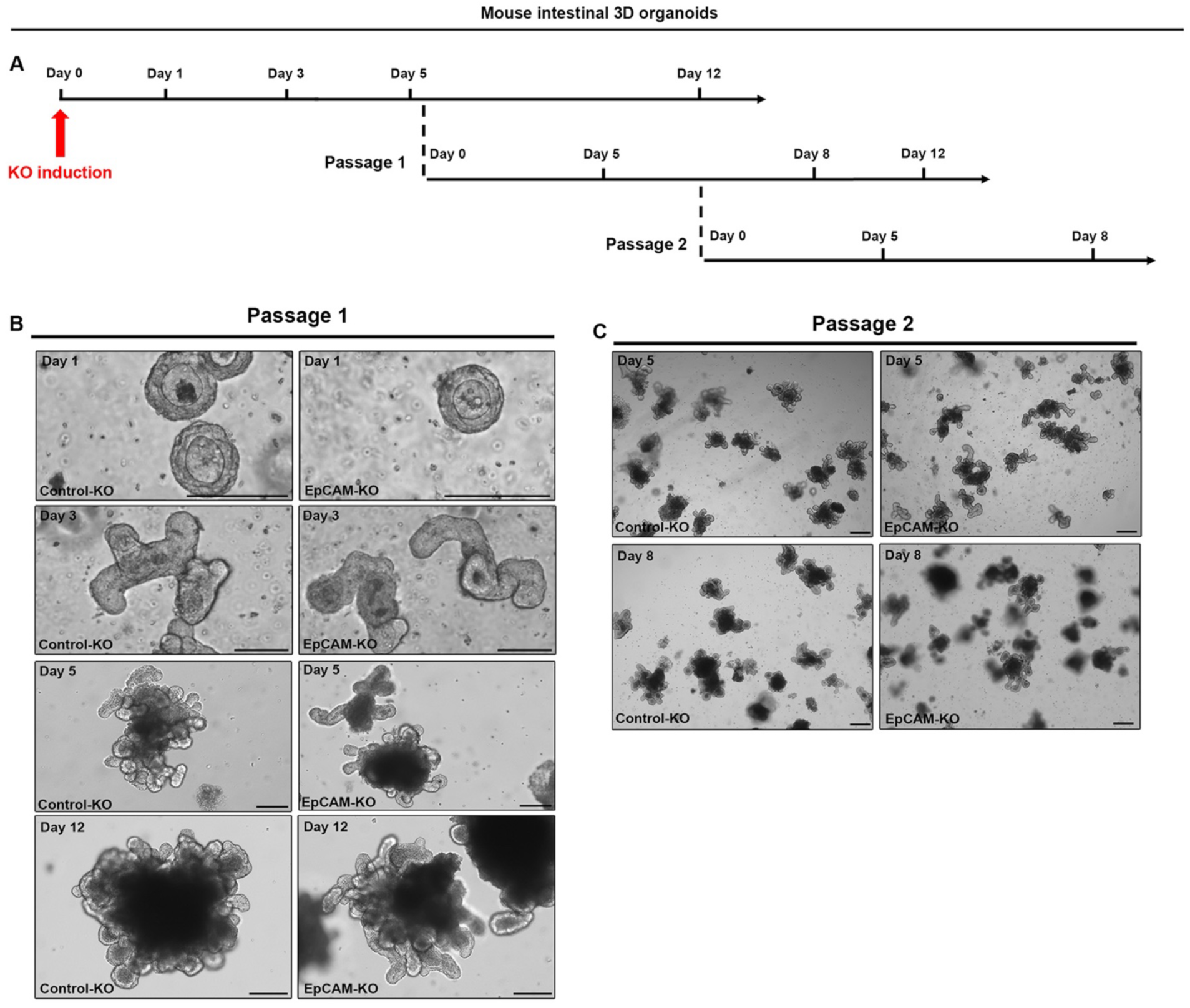
**A**, Schematic of organoid culture time courses. **B**, Phase contrast images of organoid morphology at day 1, 3, 5 or 12 of culture in control-KO or EpCAM-KO organoid lines after one post-induction passage. Scale bar, 100μm. **C**, Phase contrast images of organoid morphology at day 5 or 8 of culture in control-KO or EpCAM-KO organoid lines after two post-induction passages. Scale bar, 100μm.

**Supplementary Figure 3:**
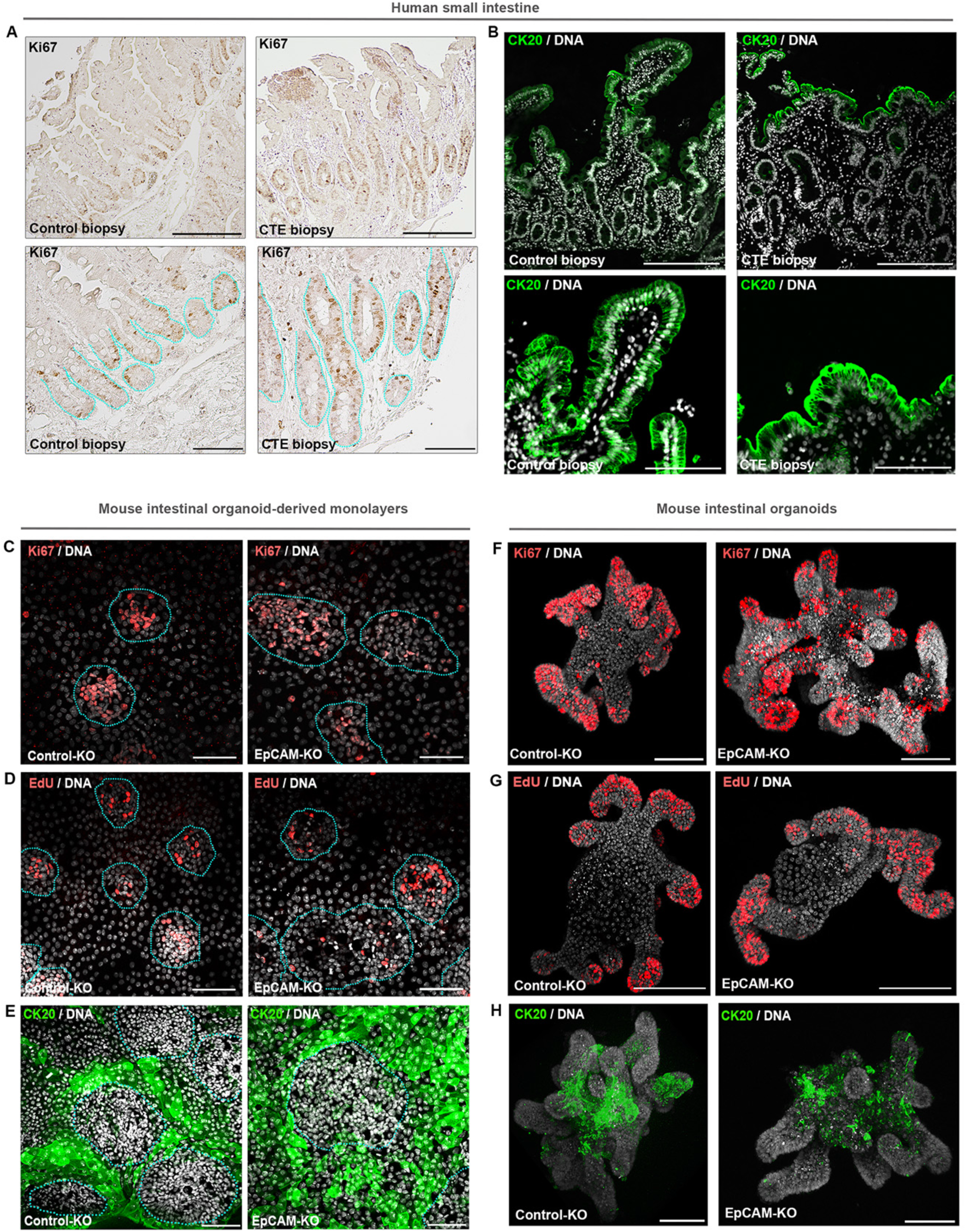
**A,** Immunohistological analysis of Ki67 (brown) localization in control and CTE biopsies. Blue dotted lines delimit the crypt compartments. Scale bars, upper panel 150μm, lower panel 50 μm. **B**, Confocal microscopy analysis of CK20 (green) distribution along the crypt-villus axis of control and CTE patients. Nuclei are shown in grey. Scale bars, upper panel 200μm, lower panel 100 μm. **C-E**, Confocal microscopy analysis of Ki67 (**C**, red), EdU (**D**, red) and CK20 (**E**, green) together with DNA (grey) in control-KO and EpCAM-KO organoid-derived monolayers. Blue dotted lines delimit the crypt-like compartments. Scale bars, **C-D** 50μm, **E**100μm. **F-H**, Confocal microscopy analysis of Ki67 (red) (**F**), EdU (red) (**G**) and CK20 (green) (**H**) in control-KO and EpCAM-KO organoids. Scale bar, 80μm.

**Supplementary Figure 4:**
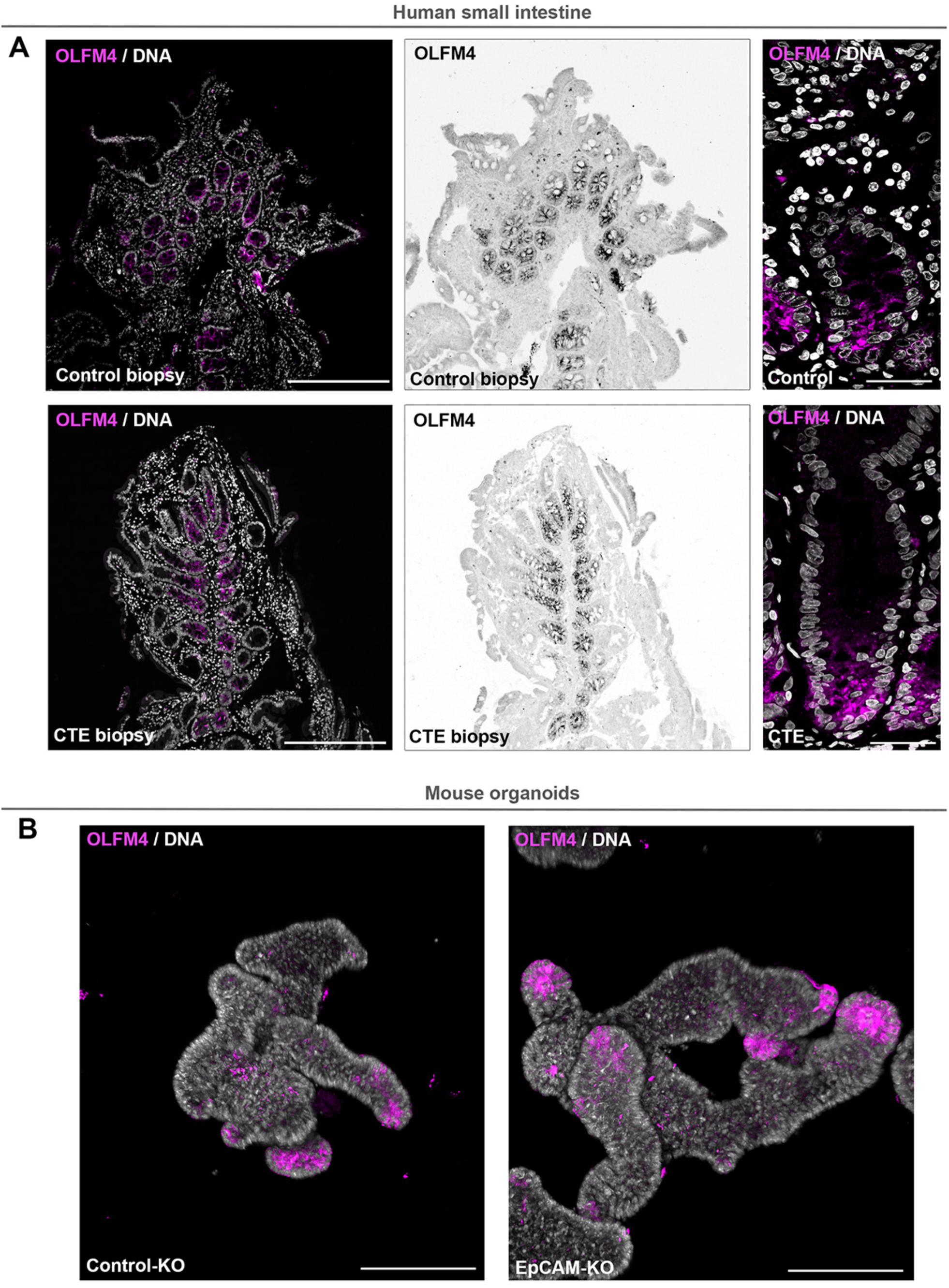
**A,** Confocal microscopy analysis of the distribution of OLFM4 (magenta) in crypts of control and EpCAM-mutated CTE patients. Scale bars, left panels 200μm, right panel 100 μm. **B**, Confocal microscopy analysis of OLFM4 (magenta) in control-KO and EpCAM-KO organoids. Scale bar, 100μm.

**Supplementary Figure 5:**
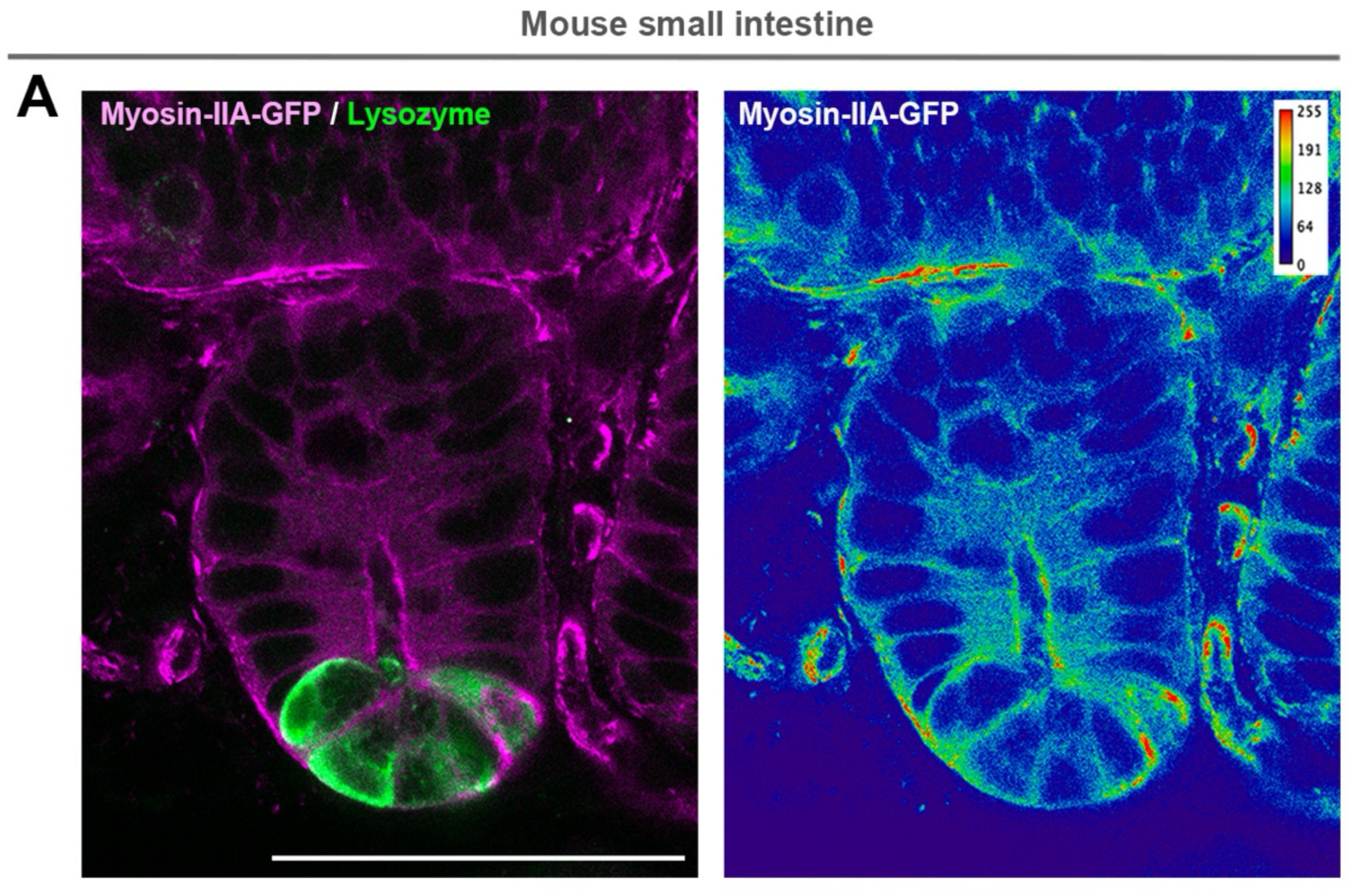
**A**, Confocal analysis of the localization of myosin-IIA (magenta) and lysozyme (green) in mouse jejunum crypts. Color-coding of myosin-II-A-GFP signal intensity with Physics LUT ImageJ is shown. Calibration bar is added as a right insert. Scale bar, 50μm.

**Supplementary Figure 6:**
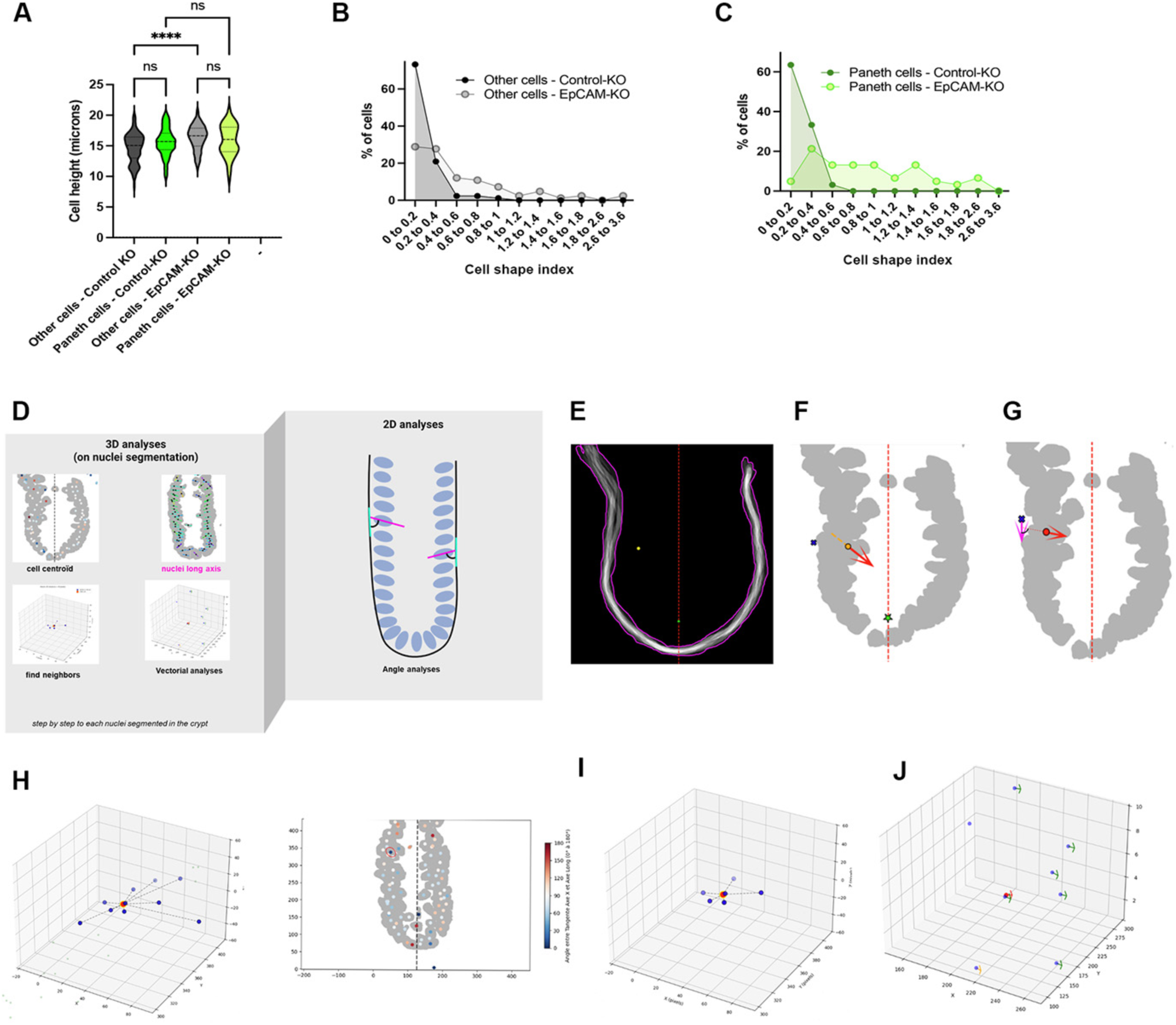
**A**, Statistical analysis of cell heights in control-KO and EpCAM-KO crypts. Mean cell height of control-KO Paneth cells = 15.59±0.29μm (mean±S.E.M), control-KO other cells= 14.67±0.26μm, EpCAM-KO Paneth cells = 15.9±0.33μm, EpCAM-KO other cells = 16.37±0.22μm. ****p<0.0001. n (control-KO Paneth cells) = 63 cells, n (control-KO other cells) = 86 cells, n (EpCAM-KO Paneth cells) = 61 cells, n (EpCAM-KO other cells) = 84 cells. One-way ANOVA, Tukey’s multiple tests, ****p<0.0001. **B**, Distribution of cell shape indexes of other crypt cells in control-KO and EpCAM-KO crypts. Cell shape indexes from 0 to 0.2: control-KO = 73.25%, EpCAM-KO = 28.91%, from 0.2 to 0.4: control-KO = 20.93%, EpCAM-KO = 27.71%, from 0.4 to 0.6: control-KO = 2.33%, EpCAM-KO = 12.05%, from 0.6 to 0.8: control-KO = 2.33%, EpCAM-KO = 10.84%, from 0.8 to 1: control-KO = 1.16%, EpCAM-KO = 7.24%, from 1 to 3.6: control-KO = 0%, EpCAM-KO = 13.25%. **C**, Distribution of cell shape indexes of Paneth cells in control-KO and EpCAM-KO crypts. Cell shape indexes from 0 to 0.2: control-KO = 63.5%, EpCAM-KO = 4.92%, from 0.2 to 0.4: control-KO = 33.33%, EpCAM-KO = 21.32%, from 0.4 to 0.6: control-KO = 3.17%, EpCAM-KO = 13.11%, from 0.6 to 0.8: control-KO = 0%, EpCAM-KO = 13.11%, from 0.8 to 1: control-KO = 0%, EpCAM-KO = 13.11%, from 1 to 3.6: control-KO = 0%, EpCAM-KO = 34.43%. **D,** 3D analysis to extract cell centroids, nuclear long axes, neighbor relationships, and perform vectorial analyses, enabling 2D projection for angle measurements within segmented crypts. **E,** Crypt contour used to define local tangents for each nucleus. Yellow dot, nucleus shown as an example; green star, reference point for angle calculation; red line, reference axis for orientation of the nuclear long axis. **F.** 2D projection of nuclei (grey). Red arrow, nuclear long axis defined in 3D; blue cross, anchoring point of the crypt contour tangent at the nucleus position. **G.** Angle measurement based on 3D data projected into 2D. Red arrow, local tangent to the crypt contour at the nucleus position. **H.** Neighbor identification based on cell centroids. Circled nucleus (red), target nucleus shown as an example. **I.** Distance threshold applied to refine neighbor detection based on neighbor matching from 3D-segmented nuclei.

**Supplementary Figure 7:**
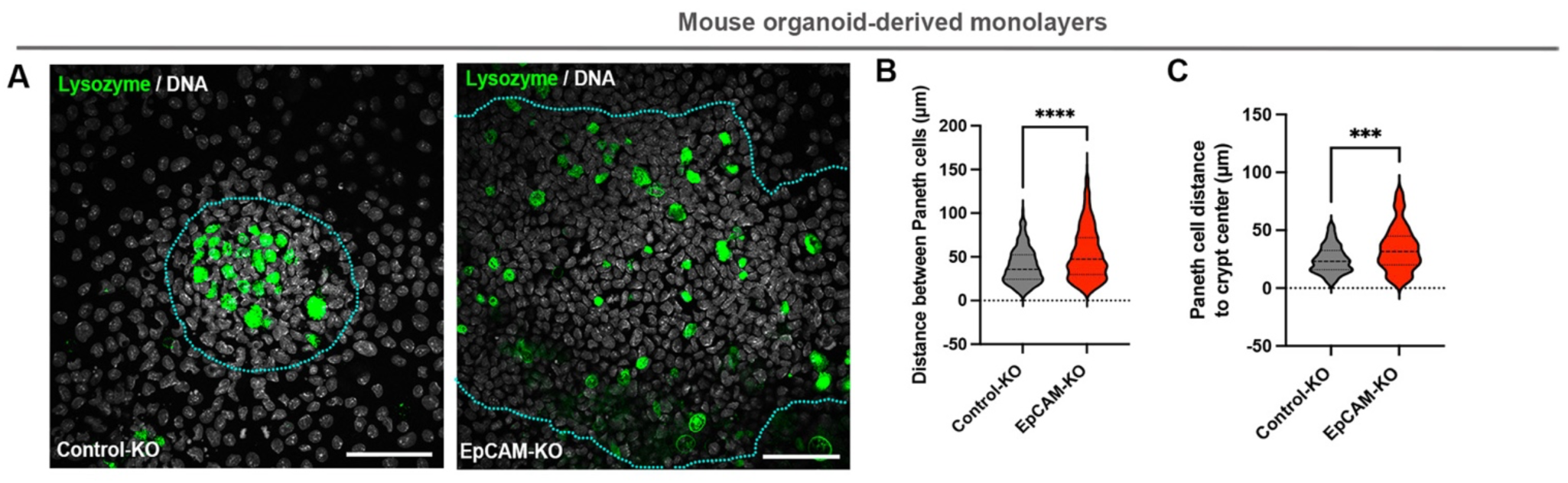
**A**, Confocal microscopy analysis of lysozyme (green) together with DNA (grey) in control-KO and EpCAM-KO organoid-derived monolayers. Blue dotted lines delimit the crypt-like compartments. Scale bar, 50μm. **B**, Statistical analysis of the mean distance between Paneth cells in organoid-derived monolayers. Distance in control-KO monolayers = 39.54±1.12μm (mean±S.E.M), in EpCAM-KO monolayers = 53.39±1.85μm. Mann-Whitney test, ****p<0.0001. n (control-KO) = 311 Paneth cells, n (EpCAM-KO) = 614 Paneth cells. **C**, Statistical analysis of the mean distance between Paneth cells and the crypt center. Distance in control-KO monolayers = 24.85±1.30μm (mean±S.E.M), in EpCAM-KO monolayers = 34.23±1.81μm. Mann-Whitney test, ***p=0.0006. n (control-KO) = 80 Paneth cells, n (EpCAM-KO) = 113 Paneth cells.

## REFERENCES

1. Petridou, N.I., Spiro, Z., and Heisenberg, C.P. (2017). Multiscale force sensing in development. Nat Cell Biol 19, 581–588. 10.1038/ncb3524.

2. Messal, H.A., Alt, S., Ferreira, R.M.M., Gribben, C., Wang, V.M., Cotoi, C.G., Salbreux, G., and Behrens, A. (2019). Tissue curvature and apicobasal mechanical tension imbalance instruct cancer morphogenesis. Nature 566, 126–130. 10.1038/s41586-019-0891-2.

3. Zinner, M., Lukonin, I., and Liberali, P. (2020). Design principles of tissue organisation: How single cells coordinate across scales. Curr Opin Cell Biol 67, 37–45. 10.1016/j.ceb.2020.07.004.

4. Sato, T., and Clevers, H. (2013). Growing self-organizing mini-guts from a single intestinal stem cell: mechanism and applications. Science 340, 1190–1194. 10.1126/science.1234852.

5. Yang, Q., Xue, S.L., Chan, C.J., Rempfler, M., Vischi, D., Maurer-Gutierrez, F., Hiiragi, T., Hannezo, E., and Liberali, P. (2021). Cell fate coordinates mechano-osmotic forces in intestinal crypt formation. Nat Cell Biol 23, 733–744. 10.1038/s41556-021-00700-2.

6. Perez-Gonzalez, C., Ceada, G., Matejcic, M., and Trepat, X. (2022). Digesting the mechanobiology of the intestinal epithelium. Curr Opin Genet Dev 72, 82–90. 10.1016/j.gde.2021.10.005.

7. Houtekamer, R.M., van der Net, M.C., Maurice, M.M., and Gloerich, M. (2022). Mechanical forces directing intestinal form and function. Curr Biol 32, R791–R805. 10.1016/j.cub.2022.05.041.

8. Barker, N. (2014). Adult intestinal stem cells: critical drivers of epithelial homeostasis and regeneration. Nat Rev Mol Cell Biol 15, 19–33. 10.1038/nrm3721.

9. van der Flier, L.G., and Clevers, H. (2009). Stem cells, self-renewal, and differentiation in the intestinal epithelium. Annu Rev Physiol 71, 241–260. 10.1146/annurev.physiol.010908.163145.

10. Guevara-Garcia, A., Soleilhac, M., Minc, N., and Delacour, D. (2023). Regulation and functions of cell division in the intestinal tissue. Semin Cell Dev Biol 150–151, 3-14. 10.1016/j.semcdb.2023.01.004.

11. Maghzal, N., Kayali, H.A., Rohani, N., Kajava, A.V., and Fagotto, F. (2013). EpCAM controls actomyosin contractility and cell adhesion by direct inhibition of PKC. Dev Cell 27, 263–277. 10.1016/j.devcel.2013.10.003.

12. Salomon, J., Gaston, C., Magescas, J., Duvauchelle, B., Canioni, D., Sengmanivong, L., Mayeux, A., Michaux, G., Campeotto, F., Lemale, J., et al. (2017). Contractile forces at tricellular contacts modulate epithelial organization and monolayer integrity. Nat Commun 8, 13998. 10.1038/ncomms13998.

13. Gaston, C., De Beco, S., Doss, B., Pan, M., Gauquelin, E., D’Alessandro, J., Lim, C.T., Ladoux, B., and Delacour, D. (2021). EpCAM promotes endosomal modulation of the cortical RhoA zone for epithelial organization. Nat Commun 12, 2226. 10.1038/s41467-021-22482-9.

14. Ladwein, M., Pape, U.F., Schmidt, D.S., Schnolzer, M., Fiedler, S., Langbein, L., Franke, W.W., Moldenhauer, G., and Zoller, M. (2005). The cell-cell adhesion molecule EpCAM interacts directly with the tight junction protein claudin-7. Exp Cell Res 309, 345–357. 10.1016/j.yexcr.2005.06.013.

15. Wu, S.K., and Yap, A.S. (2013). Patterns in space: coordinating adhesion and actomyosin contractility at E-cadherin junctions. Cell Commun Adhes 20, 201–212. 10.3109/15419061.2013.856889.

16. Trzpis, M., McLaughlin, P.M., de Leij, L.M., and Harmsen, M.C. (2007). Epithelial cell adhesion molecule: more than a carcinoma marker and adhesion molecule. Am J Pathol 171, 386–395.

17. Sivagnanam, M., Mueller, J.L., Lee, H., Chen, Z., Nelson, S.F., Turner, D., Zlotkin, S.H., Pencharz, P.B., Ngan, B.Y., Libiger, O., et al. (2008). Identification of EpCAM as the gene for congenital tufting enteropathy. Gastroenterology 135, 429–437.

18. Salomon, J., Goulet, O., Canioni, D., Brousse, N., Lemale, J., Tounian, P., Coulomb, A., Marinier, E., Hugot, J.P., Ruemmele, F., et al. (2014). Genetic characterization of congenital tufting enteropathy: epcam associated phenotype and involvement of SPINT2 in the syndromic form. Hum Genet 133, 299–310. 10.1007/s00439-013-1380-6.

19. Patey, N., Scoazec, J.Y., Cuenod-Jabri, B., Canioni, D., Kedinger, M., Goulet, O., and Brousse, N. (1997). Distribution of cell adhesion molecules in infants with intestinal epithelial dysplasia (tufting enteropathy). Gastroenterology 113, 833–843.

20. Goulet, O., Salomon, J., Ruemmele, F., de Serres, N.P., and Brousse, N. (2007). Intestinal epithelial dysplasia (tufting enteropathy). Orphanet J Rare Dis 2, 20. 10.1186/1750-1172-2-20.

21. Delacour, D., Salomon, J., Robine, S., and Louvard, D. (2016). Plasticity of the brush border - the yin and yang of intestinal homeostasis. Nat Rev Gastroenterol Hepatol. 10.1038/nrgastro.2016.5.

22. Bosveld, F., Wang, Z., and Bellaiche, Y. (2018). Tricellular junctions: a hot corner of epithelial biology. Curr Opin Cell Biol 54, 80–88. 10.1016/j.ceb.2018.05.002.

23. Cheng, H., Bjerknes, M., Amar, J., and Gardiner, G. (1986). Crypt production in normal and diseased human colonic epithelium. Anat Rec 216, 44–48. 10.1002/ar.1092160108.

24. Serra, D., Mayr, U., Boni, A., Lukonin, I., Rempfler, M., Challet Meylan, L., Stadler, M.B., Strnad, P., Papasaikas, P., Vischi, D., et al. (2019). Self-organization and symmetry breaking in intestinal organoid development. Nature 569, 66–72. 10.1038/s41586-019-1146-y.

25. Sumigray, K.D., Terwilliger, M., and Lechler, T. (2018). Morphogenesis and Compartmentalization of the Intestinal Crypt. Dev Cell 45, 183–197 e185. 10.1016/j.devcel.2018.03.024.

26. Perez-Gonzalez, C., Ceada, G., Greco, F., Matejcic, M., Gomez-Gonzalez, M., Castro, N., Menendez, A., Kale, S., Krndija, D., Clark, A.G., et al. (2021). Mechanical compartmentalization of the intestinal organoid enables crypt folding and collective cell migration. Nat Cell Biol 23, 745–757. 10.1038/s41556-021-00699-6.

27. Tallapragada, N.P., Cambra, H.M., Wald, T., Keough Jalbert, S., Abraham, D.M., Klein, O.D., and Klein, A.M. (2021). Inflation-collapse dynamics drive patterning and morphogenesis in intestinal organoids. Cell Stem Cell 28, 1516–1532 e1514. 10.1016/j.stem.2021.04.002.

28. Huang, L., Yang, Y., Yang, F., Liu, S., Zhu, Z., Lei, Z., and Guo, J. (2018). Functions of EpCAM in physiological processes and diseases (Review). Int J Mol Med 42, 1771–1785. 10.3892/ijmm.2018.3764.

29. Sui, L., Alt, S., Weigert, M., Dye, N., Eaton, S., Jug, F., Myers, E.W., Julicher, F., Salbreux, G., and Dahmann, C. (2018). Differential lateral and basal tension drive folding of Drosophila wing discs through two distinct mechanisms. Nat Commun 9, 4620. 10.1038/s41467-018-06497-3.

30. Xi, W., Saleh, J., Yamada, A., Tomba, C., Mercier, B., Janel, S., Dang, T., Soleilhac, M., Djemat, A., Wu, H., et al. (2022). Modulation of designer biomimetic matrices for optimized differentiated intestinal epithelial cultures. Biomaterials 282, 121380. 10.1016/j.biomaterials.2022.121380.

31. Clevers, H.C., and Bevins, C.L. (2013). Paneth cells: maestros of the small intestinal crypts. Annu Rev Physiol 75, 289–311. 10.1146/annurev-physiol-030212-183744.

32. Mei, X., Gu, M., and Li, M. (2020). Plasticity of Paneth cells and their ability to regulate intestinal stem cells. Stem Cell Res Ther 11, 349. 10.1186/s13287-020-01857-7.

33. Langlands, A.J., Almet, A.A., Appleton, P.L., Newton, I.P., Osborne, J.M., and Nathke, I.S. (2016). Paneth Cell-Rich Regions Separated by a Cluster of Lgr5+ Cells Initiate Crypt Fission in the Intestinal Stem Cell Niche. PLoS Biol 14, e1002491. 10.1371/journal.pbio.1002491.

34. Totafurno, J., Bjerknes, M., and Cheng, H. (1987). The crypt cycle. Crypt and villus production in the adult intestinal epithelium. Biophys J 52, 279–294. 10.1016/S0006-3495(87)83215-0.

35. Dudhwala, Z.M., Hammond, P.D., Howarth, G.S., and Cummins, A.G. (2020). Intestinal stem cells promote crypt fission during postnatal growth of the small intestine. BMJ Open Gastroenterol 7. 10.1136/bmjgast-2020-000388.

36. Bruens, L., Ellenbroek, S.I.J., van Rheenen, J., and Snippert, H.J. (2017). In Vivo Imaging Reveals Existence of Crypt Fission and Fusion in Adult Mouse Intestine. Gastroenterology 153, 674–677 e673. 10.1053/j.gastro.2017.05.019.

37. Boman, B.M., and Fields, J.Z. (2013). An APC:WNT Counter-Current-Like Mechanism Regulates Cell Division Along the Human Colonic Crypt Axis: A Mechanism That Explains How APC Mutations Induce Proliferative Abnormalities That Drive Colon Cancer Development. Front Oncol 3, 244. 10.3389/fonc.2013.00244.

38. Rubio, C.A., and Schmidt, P.T. (2018). Are Non-dysplastic Crypts with Corrupted Shapes the Initial Recordable Histological Event in the Development of Sporadic Conventional Adenomas? Anticancer Res 38, 5315–5320. 10.21873/anticanres.12858.

39. Dekaney, C.M., Gulati, A.S., Garrison, A.P., Helmrath, M.A., and Henning, S.J. (2009). Regeneration of intestinal stem/progenitor cells following doxorubicin treatment of mice. Am J Physiol Gastrointest Liver Physiol 297, G461–470. 10.1152/ajpgi.90446.2008.

40. Cairnie, A.B., and Millen, B.H. (1975). Fission of crypts in the small intestine of the irradiated mouse. Cell Tissue Kinet 8, 189–196. 10.1111/j.1365-2184.1975.tb01219.x.

41. Huang, X.T., Li, T., Li, T., Xing, S., Tian, J.Z., Ding, Y.F., Cai, S.L., Yang, Y.S., Wood, C., Yang, J.S., and Yang, W.J. (2022). Embryogenic stem cell-derived intestinal crypt fission directs de novo crypt genesis. Cell Rep 41, 111796. 10.1016/j.celrep.2022.111796.

42. Dekaney, C.M., Fong, J.J., Rigby, R.J., Lund, P.K., Henning, S.J., and Helmrath, M.A. (2007). Expansion of intestinal stem cells associated with long-term adaptation following ileocecal resection in mice. Am J Physiol Gastrointest Liver Physiol 293, G1013–1022. 10.1152/ajpgi.00218.2007.

43. Litvinov, S.V., Balzar, M., Winter, M.J., Bakker, H.A., Briaire-de Bruijn, I.H., Prins, F., Fleuren, G.J., and Warnaar, S.O. (1997). Epithelial cell adhesion molecule (Ep-CAM) modulates cell-cell interactions mediated by classic cadherins. J Cell Biol 139, 1337–1348.

44. Maetzel, D., Denzel, S., Mack, B., Canis, M., Went, P., Benk, M., Kieu, C., Papior, P., Baeuerle, P.A., Munz, M., and Gires, O. (2009). Nuclear signalling by tumour-associated antigen EpCAM. Nat Cell Biol 11, 162–171. 10.1038/ncb1824.

45. Slanchev, K., Carney, T.J., Stemmler, M.P., Koschorz, B., Amsterdam, A., Schwarz, H., and Hammerschmidt, M. (2009). The epithelial cell adhesion molecule EpCAM is required for epithelial morphogenesis and integrity during zebrafish epiboly and skin development. PLoS Genet 5, e1000563. 10.1371/journal.pgen.1000563.

46. Guerra, E., Lattanzio, R., La Sorda, R., Dini, F., Tiboni, G.M., Piantelli, M., and Alberti, S. (2012). mTrop1/Epcam Knockout Mice Develop Congenital Tufting Enteropathy through Dysregulation of Intestinal E-cadherin/beta-catenin. PLoS One 7, e49302.

47. Rauzi, M., Verant, P., Lecuit, T., and Lenne, P.F. (2008). Nature and anisotropy of cortical forces orienting Drosophila tissue morphogenesis. Nat Cell Biol 10, 1401–1410. 10.1038/ncb1798.

48. Fernandez-Gonzalez, R., and Zallen, J.A. (2009). Cell mechanics and feedback regulation of actomyosin networks. Sci Signal 2, pe78. 10.1126/scisignal.2101pe78.

49. Pentinmikko, N., Lozano, R., Scharaw, S., Andersson, S., Englund, J.I., Castillo-Azofeifa, D., Gallagher, A., Broberg, M., Song, K.Y., Sola Carvajal, A., et al. (2022). Cellular shape reinforces niche to stem cell signaling in the small intestine. Sci Adv 8, eabm1847. 10.1126/sciadv.abm1847.

50. Shyer, A.E., Huycke, T.R., Lee, C., Mahadevan, L., and Tabin, C.J. (2015). Bending gradients: how the intestinal stem cell gets its home. Cell 161, 569–580. 10.1016/j.cell.2015.03.041.

51. Quintero, M., and Samuelson, L.C. (2025). Paneth Cells: Dispensable yet Irreplaceable for the Intestinal Stem Cell Niche. Cell Mol Gastroenterol Hepatol 19, 101443. 10.1016/j.jcmgh.2024.101443.

52. Yavitt, F.M., Khang, A., Bera, K., McNally, D.L., Blatchley, M.R., Gallagher, A.P., Klein, O.D., Castillo-Azofeifa, D., Dempsey, P.J., and Anseth, K.S. (2025). Engineered epithelial curvature controls Paneth cell localization in intestinal organoids. Cell Biomater 1. 10.1016/j.celbio.2025.100046.

53. Cummins, A.G., Jones, B.J., and Thompson, F.M. (2006). Postnatal epithelial growth of the small intestine in the rat occurs by both crypt fission and crypt hyperplasia. Dig Dis Sci 51, 718–723. 10.1007/s10620-006-3197-9.

54. Slupecka M., W.J., Pierzynowski S. (2010). Crypt fission contributes to postnatal epithelial growth of the small intestine in pigs. Livestock Science 133, 34–37.

55. Baker, A.M., Cereser, B., Melton, S., Fletcher, A.G., Rodriguez-Justo, M., Tadrous, P.J., Humphries, A., Elia, G., McDonald, S.A., Wright, N.A., et al. (2014). Quantification of crypt and stem cell evolution in the normal and neoplastic human colon. Cell Rep 8, 940–947. 10.1016/j.celrep.2014.07.019.

56. Cummins, A.G., Catto-Smith, A.G., Cameron, D.J., Couper, R.T., Davidson, G.P., Day, A.S., Hammond, P.D., Moore, D.J., and Thompson, F.M. (2008). Crypt fission peaks early during infancy and crypt hyperplasia broadly peaks during infancy and childhood in the small intestine of humans. J Pediatr Gastroenterol Nutr 47, 153–157. 10.1097/MPG.0b013e3181604d27.

57. Rubio, C.A., Schmidt, P.T., Lang-Schwarz, C., and Vieth, M. (2022). Branching crypts in inflammatory bowel disease revisited. J Gastroenterol Hepatol 37, 440–445. 10.1111/jgh.15734.

58. Quyn, A.J., Appleton, P.L., Carey, F.A., Steele, R.J., Barker, N., Clevers, H., Ridgway, R.A., Sansom, O.J., and Nathke, I.S. (2010). Spindle orientation bias in gut epithelial stem cell compartments is lost in precancerous tissue. Cell Stem Cell 6, 175–181. 10.1016/j.stem.2009.12.007.

59. Pin, C., Parker, A., Gunning, A.P., Ohta, Y., Johnson, I.T., Carding, S.R., and Sato, T. (2015). An individual based computational model of intestinal crypt fission and its application to predicting unrestrictive growth of the intestinal epithelium. Integr Biol (Camb) 7, 213–228. 10.1039/c4ib00236a.

60. Wong, W.M., Mandir, N., Goodlad, R.A., Wong, B.C., Garcia, S.B., Lam, S.K., and Wright, N.A. (2002). Histogenesis of human colorectal adenomas and hyperplastic polyps: the role of cell proliferation and crypt fission. Gut 50, 212–217. 10.1136/gut.50.2.212.

61. Humphries, A., and Wright, N.A. (2008). Colonic crypt organization and tumorigenesis. Nat Rev Cancer 8, 415–424. 10.1038/nrc2392.

62. Wasan, H.S., Park, H.S., Liu, K.C., Mandir, N.K., Winnett, A., Sasieni, P., Bodmer, W.F., Goodlad, R.A., and Wright, N.A. (1998). APC in the regulation of intestinal crypt fission. J Pathol 185, 246–255. 10.1002/(SICI)1096-9896(199807)185:3<246::AID-PATH90>3.0.CO;2-8.

63. Liu, Y., Wang, Y., Sun, S., Chen, Z., Xiang, S., Ding, Z., Huang, Z., and Zhang, B. (2022). Understanding the versatile roles and applications of EpCAM in cancers: from bench to bedside. Exp Hematol Oncol 11, 97. 10.1186/s40164-022-00352-4.

64. Patriarca, C., Macchi, R.M., Marschner, A.K., and Mellstedt, H. (2012). Epithelial cell adhesion molecule expression (CD326) in cancer: a short review. Cancer Treat Rev 38, 68–75. 10.1016/j.ctrv.2011.04.002.

65. Kovacs, M.E., Papp, J., Szentirmay, Z., Otto, S., and Olah, E. (2009). Deletions removing the last exon of TACSTD1 constitute a distinct class of mutations predisposing to Lynch syndrome. Hum Mutat 30, 197–203. 10.1002/humu.20942.

66. Ligtenberg, M.J., Kuiper, R.P., Chan, T.L., Goossens, M., Hebeda, K.M., Voorendt, M., Lee, T.Y., Bodmer, D., Hoenselaar, E., Hendriks-Cornelissen, S.J., et al. (2009). Heritable somatic methylation and inactivation of MSH2 in families with Lynch syndrome due to deletion of the 3’ exons of TACSTD1. Nat Genet 41, 112–117. 10.1038/ng.283.

67. Lynch, H.T., Snyder, C.L., Shaw, T.G., Heinen, C.D., and Hitchins, M.P. (2015). Milestones of Lynch syndrome: 1895-2015. Nat Rev Cancer 15, 181–194. 10.1038/nrc3878.

68. Krndija, D., El Marjou, F., Guirao, B., Richon, S., Leroy, O., Bellaiche, Y., Hannezo, E., and Matic Vignjevic, D. (2019). Active cell migration is critical for steady-state epithelial turnover in the gut. Science 365, 705–710. 10.1126/science.aau3429.

69. Saleh, J., Fardin, M.A., Barai, A., Soleilhac, M., Frenoy, O., Gaston, C., Cui, H., Dang, T., Gaudin, N., Vincent, A., et al. (2023). Length limitation of astral microtubules orients cell divisions in murine intestinal crypts. Dev Cell 58, 1519–1533 e1516. 10.1016/j.devcel.2023.06.004.

70. Barai A, S.M., Xi W, Lin SZ, Karnat M, Bazellières E, Richelme S, Berrebi D, Ruemmele F, Théry M, Rupprecht JF, Delacour D (2025). Multicellular actin star network underpins epithelial organization and connectivity. Nature Communications in press. 10.1101/2024.07.26.605277

71. Miyoshi, H., and Stappenbeck, T.S. (2013). In vitro expansion and genetic modification of gastrointestinal stem cells in spheroid culture. Nat Protoc 8, 2471–2482. 10.1038/nprot.2013.153.

72. Barai, A., Das, A., and Sen, S. (2021). Measuring microenvironment-tuned nuclear stiffness of cancer cells with atomic force microscopy. STAR Protoc 2, 100296. 10.1016/j.xpro.2021.100296.

73. Marshall, A.R., Maniou, E., Moulding, D., Greene, N.D.E., Copp, A.J., and Galea, G.L. (2022). Two-Photon Cell and Tissue Level Laser Ablation Methods to Study Morphogenetic Biomechanics. Methods Mol Biol 2438, 217–230. 10.1007/978-1-0716-2035-9_14.

